# Epigenome and transcriptome changes in *KMT2D*-related Kabuki syndrome Type 1 iPSCs, neuronal progenitors and cortical neurons

**DOI:** 10.1101/2025.02.06.636815

**Authors:** S. Cuvertino, E. Martirosian, P. Cheng, T. Garner, I. J. Donaldson, A. Jackson, A. Stevens, A. D. Sharrocks, S. J. Kimber, S. Banka

**Affiliations:** Division of Evolution, Infection and Genomics, School of Biological Sciences, Faculty of Biology, Medicine, and Health, The University of Manchester, Manchester, UK; Division of Developmental Biology & Medicine, School of Biological Sciences, Faculty of Biology, Medicine, and Health, The University of Manchester, Manchester, UK; Bioinformatics Core Facility, The University of Manchester, Manchester, UK; Manchester Centre for Genomic Medicine, St. Mary’s Hospital, Manchester University Foundation NHS Trust, Health Innovation Manchester, Manchester, UK; Division of Molecular and Cellular Function, School of Biological Sciences, Faculty of Biology, Medicine and Health, The University of Manchester, Manchester, UK; Division of Cell Matrix Biology and Regenerative Medicine, School of Biological Sciences, Faculty of Biology, Medicine, and Health, The University of Manchester, Manchester, UK

## Abstract

Kabuki syndrome type 1 (KS1) is a neurodevelopmental disorder caused by loss-of-function variants in *KMT2D* which encodes a H3K4 methyltransferase. The mechanisms underlying neurodevelopmental problems in KS1 are still largely unknown. Here, we track the epigenome and transcriptome across three stages of neuronal differentiation using patient-derived induced pluripotent stem cells (iPSCs) to gain insights into the disease mechanism of KS1. In KS1 iPSCs we detected significantly lower levels of functional *KMT2D* transcript and KMT2D protein, and lower global H3K4me1 and H3K4me2 levels. We identify loss of thousands of H3K4me1 peaks in iPSCs, neuronal progenitors (NPs) and early cortical neurons (CNs) in KS1. We show that the number of lost peaks increase as differentiation progresses. We also identify hundreds of differentially expressed genes (DEGs) in iPSCs, NPs and CNs in KS1. In contrast with the epigenomic changes, the number of DEGs decrease as differentiation progresses. Our analysis reveals significant enrichment of differentially downregulated genes in areas containing putative enhancer regions with H3K4me1 loss. We also identify a set of distinct transcription factor binding sites in differentially methylated regions and a set of DEGs related to KS1 phenotypes. We find that genes regulated by SUZ12, a subunit of Polycomb Repressive complex 2, are over-represented in KS1 DEGs at early stages of differentiation. In conclusion, we present a disease-relevant human cellular model for KS1 that provides mechanistic insights for the disorder and could be used for high throughput drug screening for KS1.

## INTRODUCTION

*KMT2D* encodes a trithorax-related histone 3 lysine 4 (H3K4) methyltransferase, which is part of the COMPASS complex that primarily binds to enhancer regions and maintains global levels of H3K4me1(1, 2). Mouse *Kmt2d* knockouts are embryonic lethal(3) and it is required for exit of mouse embryonic stem cells (ESCs) from the naive pluripotent state, but is dispensable for maintenance of cell-identity(4, 5). KMT2D is also crucial for the development of several organs including the brain(6, 7).

Rare mono-allelic *KMT2D* variants cause Kabuki syndrome (KS) Type 1 (KS1, OMIM#147920)(8, 9). KS type 2 is caused by rare hetero- or hemizygous variants in a H3K27 demethylase encoding *KDM6A* gene (OMIM#300867)(10, 11). KS1 is characterised by distinct facial dysmorphism, mild to moderate intellectual disability, developmental delay, a range of internal organ malformations (congenital heart defects, skeletal defects, cleft palate and genitourinary malformations) and systemic problems (endocrine disorders, deafness, immune defects)(12).

Current animal and cellular models for KS1 have provided valuable insights into the disease mechanisms(7, 13, 14). *Kmt2d* zebrafish morphants showing reduced hindbrain and midbrain size(6), and learning and memory impairment were reported in a *Kmt2d^+/βGeo^* murine model(7). Suppression of oxygen-responsive gene expression has been seen in induced pluripotent stem cells (iPSCs) derived from patients with nonsense *KMT2D* variants(15). Changes in locus-specific loss of gene expression were reported in KS1 iPSC models with truncating *KMT2D* variants(16). Of note, most studies have used models with null variants, even though ∼10% KS1-causing variants are missense(17). Furthermore, most studies have used single cell types, thus our knowledge about the impact of KS1-causing *KMT2D* variants at different stages of differentiation is lacking.

Here, we track the epigenomic and transcriptional panorama across three stages of neuronal differentiation using patient-derived iPSCs to gain insights into disease mechanism of KS1. We show that in iPSCs, KS1 causing variants result in loss of KMT2D function and alter the H3K4me1 and transcriptional landscapes in a correlated manner. We show that epigenomic differences between KS1 and control cells increase with differentiation progression, but the transcriptional changes become less abundant. Our analysis also reveals a set of potential transcriptional regulators and their cis-regulatory elements, differentially expressed genes (DEGs) and master regulators that likely underpin the KS1 phenotypes.

## RESULTS

### KS1 causing variants result in loss of *KMT2D* function in iPSCs

IPSCs were generated either from fibroblasts in the HipSci program (www.hipsci.org/#/cells) or in house from fibroblasts or peripheral blood mononuclear cells and used three controls and three individuals with molecularly confirmed KS1. Our cohort included one individual each with frameshift (NM_003482.4: *KMT2D*; c.5527dupA, p.(Pro1849Ter)), missense (c.16019G>A, p.(Arg5340GIn)), and nonsense (c.16180G>T, p.(Glu5394Ter)) variants (**Figure 1A** and **Supplementary Table 1**).

**Figure 1.**
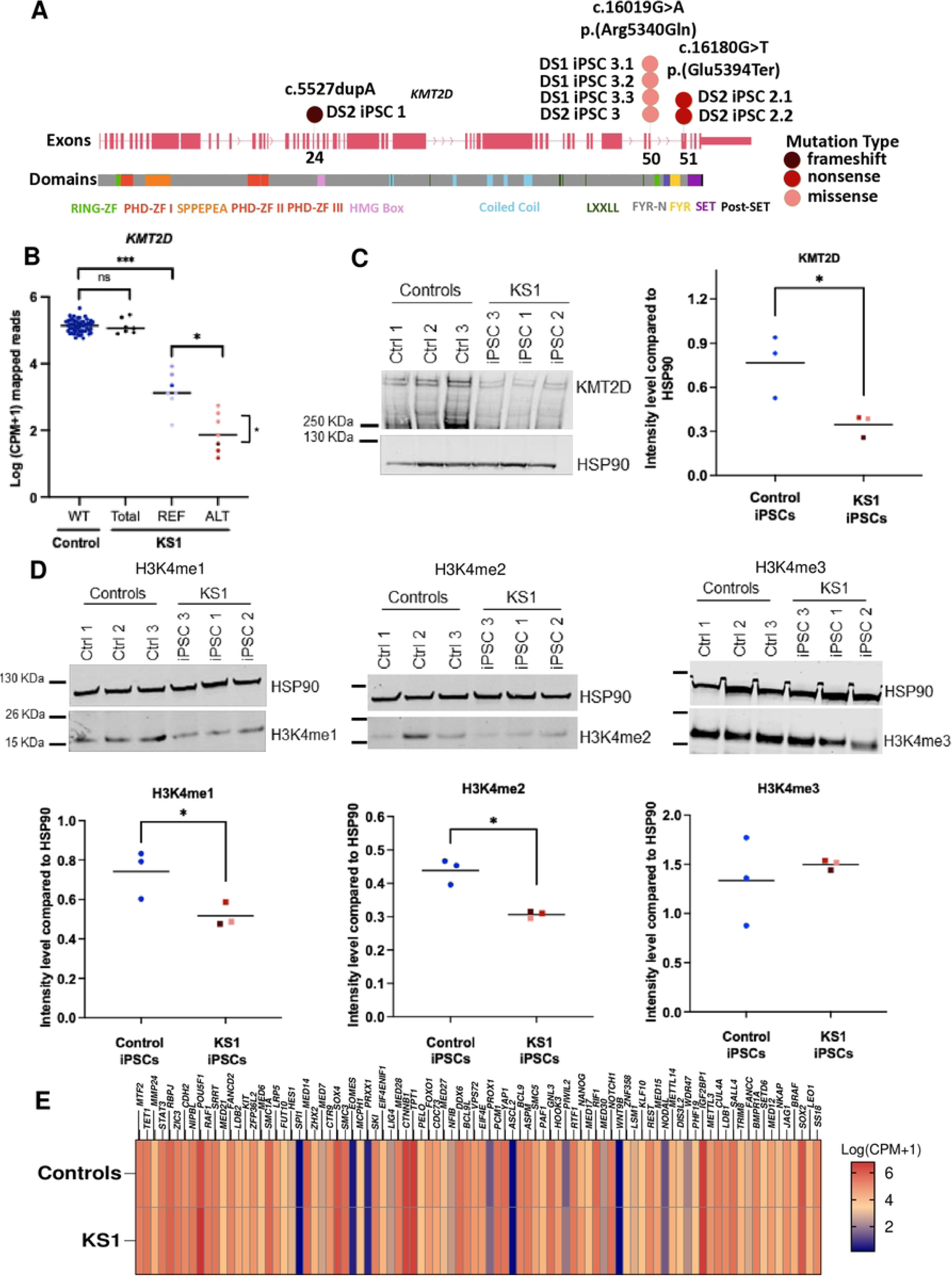
Analysis of KMT2D expression in KS1 iPSCs. **A)** Schematic representation of *KMT2D* exons and protein domains and regions with the different *KMT2D* variants highlighted in light pink, dark pink and red for missense, nonsense and frameshift mutations, respectively. (Dataset 1: DS1; Dataset 2: DS2). **B)** Log counts per million (CPM+1) mapped reads for *KMT2D* transcript detected in the integrated RNAseq analysis. REF shows reads with reference – no mutation (light blue; ***p<0.0001, *p<0.05) and ALT shows reads with mutation (pink/red). **C)** Western blot for KMT2D and scatter plot showing KMT2D quantification relative to loading control, HSP90 (n=3; * p<0.05). **D)** Representative image for Western blots for H3K4me1, H3K4me2 and H3K4me3 and scatter plots showing their quantification relative to loading control, HSP90 (n=3; *p<0.05). **E)** Log counts per million (CPM+1) mapped reads for pluripotency related transcripts detected in the integrated RNAseq analysis.

Firstly, to investigate the effect of KS1 variants at the pluripotent stem cell stage, we generated a merged iPSC RNAseq dataset of 53 control lines and 7 KS1 lines (4 replicates with the missense variant, 2 with the nonsense variant and one with the frameshift variant) (**Supplementary Figure S1**). Most disease causing *KMT2D* variants are predicted to result in haploinsufficiency but it has not been proven to the best of our knowledge(8). We, therefore, assessed the level of *KMT2D* mRNA using the merged RNAseq data. We did not observe significant difference in the total *KMT2D* mRNA level between controls and KS1 iPSCs (total *KMT2D* reads) (**Figure 1B** and **Supplementary Figure S2A**). However, quantification of reference and mutant *KMT2D* transcripts revealed that the reference *KMT2D* reads (REF) were significantly lower in all KS1 iPSCs compared to controls (**Figure 1B** and **Supplementary Figure S2B-C**). Interestingly, level of reference *KMT2D* reads (REF) in KS1 iPSCs was higher compared to reads with mutations (ALT) (**Figure 1B**). Furthermore, level of reads with missense variants was significantly higher compared to the reads with truncating variants, due to potential action of nonsense-mediated decay (**Figure 1B**).

Next, we performed Western blotting on nuclear protein extracts from iPSCs, which revealed a significant decrease in KMT2D protein level in nuclei of all KS1 iPSCs, including the one with the missense variant, compared to control iPSCs (**Figure 1C**). As KMT2D is a H3K4 methyltransferase, we quantified the total levels of H3K4me1, H3K4me2, and H3K4me3 histone marks in all three KS1 and control iPSCs by Western blotting. In all three lines, including in the one with the missense variant, we observed a significant decrease in the levels of H3K4me1 and H3K4me2 histone marks but not in H3K4me3 histone mark (**Figure 1D**). Next, we examined the DepMap portal (https://depmap.org/portal/) for the levels of 42 different histone marks in cancer cell lines with *KMT2D* mutations and compared to cell lines without *KMT2D* damaging mutations (**Supplementary Figure S3**). We found H3K4me1 and H2K4me2 levels to be most significantly reduced in cell lines with *KMT2D* mutations compared to cell lines without *KMT2D* mutations.

Next, we evaluated if loss of KMT2D function led to any changes in the molecular pluripotency state of KS1 iPSCs. In the merged RNAseq dataset, we did not find significant differences in the expression of key genes related to pluripotency in KS1 iPSCs compared to control iPSCs (**Figure 1E**). We validated these results by RTq-PCR and immunofluorescence staining for SOX2, NANOG and OCT4 (**Supplementary Figure S4**).

Overall, these data show that KS1-causing heterozygous *KMT2D* variants, including missense variants, are associated with loss of function that is reflected in significant decrease in levels of H3K4me1/me2. The loss of function resulting from KS1-causing heterozygous *KMT2D* variants does not result in major changes in pluripotency-associated gene expression.

### H3K4me1 landscape is dysregulated in KS1 iPSCs

KMT2D is a major H3K4 mono-methyltransferase essential for enhancer activation during cell differentiation(1, 3). Since we observed a significant global decrease in H3K4me1 in our cell lines (**Figure 1D**), we decided to investigate the H3K4me1 landscape in KS1 in more detail. As limited studies have been conducted on KS1 animal or cellular models with missense variants, we performed H3K4me1 ChIPseq in one KS1 (c.16019G>A, p.(Arg5340GIn)) and one control iPSC line, both in triplicate. We detected 9,309 out of 205,973 peak sites to be significantly differentially methylated (FDR<0.05; -1>log_2_FC>1). Of these, 8,146 sites (87.5% of all differentially methylated peak sites) showed loss, and 1,163 sites showed gain in the H3K4me1 peak signal (**Figure 2A-C**). H3K4me1 peak site losses were enriched in cell-type independent EpiMap annotated enhancer regions (fisher’s test; p-value < 2.2e^-16^) (**Figure 2C** and **Supplementary Figure S5A** and **S6**).

**Figure 2.**
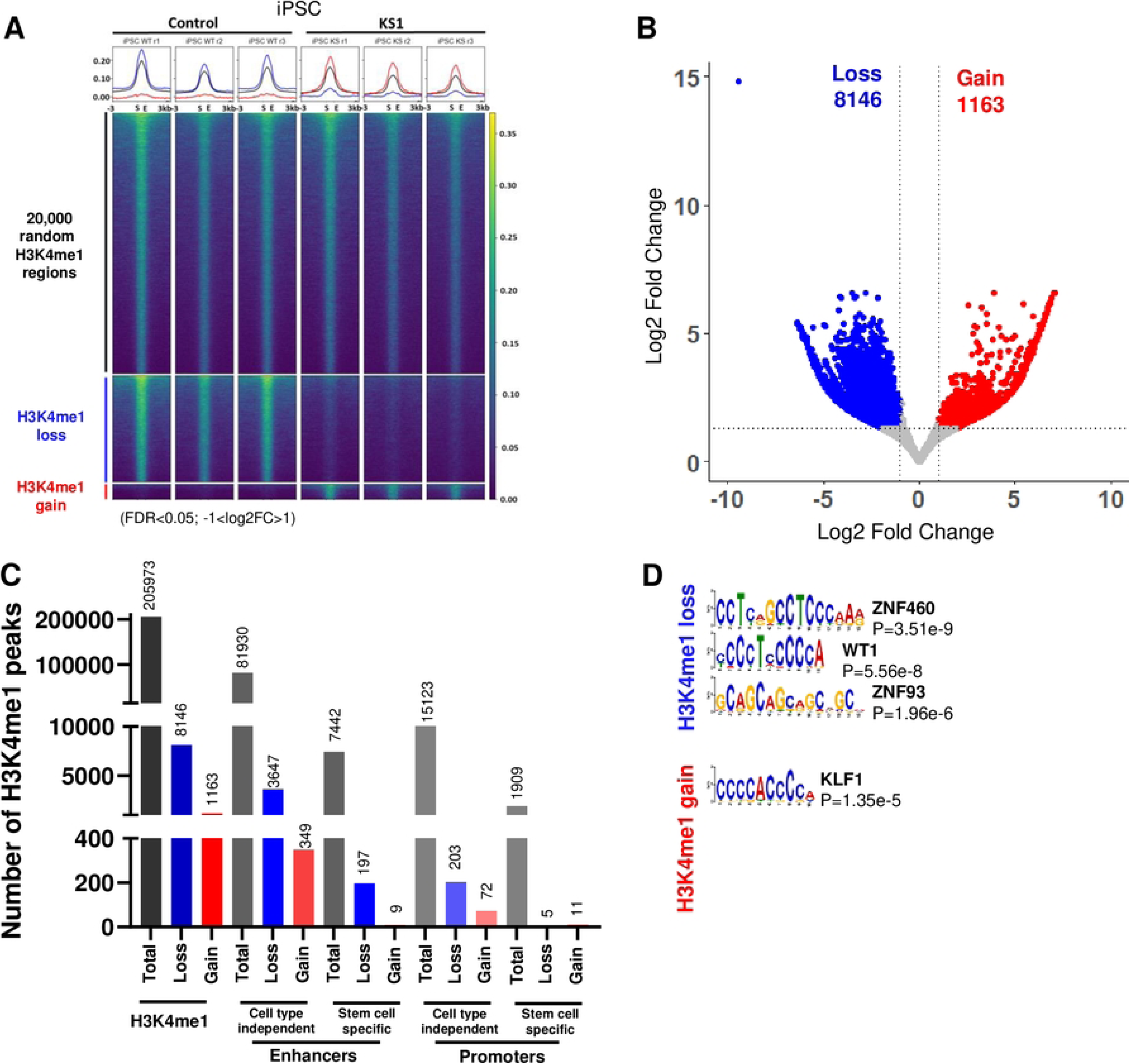
Loss of H3K4me1 in KS1 iPSCs. **A)** Heatmap of H3K4me1 peaks (subset of consensus regions 20,000 out of 205,973; black), H3K4me1 loss regions (n=8146; blue), and H3K4me1 gain regions (n=1163; red). **B)** Volcano plot showing loss (blue) and gain (red) in H3K4me1. **C)** Bar graph showing number of H3K4me1 peaks in relation to enhancer and promoter regions for total, loss and gain in mono-methylation. **D)** Top TF motifs predicted in differential H3K4me1 peaks (loss and gain of H3K4me1 in binding regions) between KS1 and control.

To identify transcription factor binding sites predicted to be affected by changes in H3K4me1 levels, we performed motif analysis on significantly differentially methylated peak sites. This showed these sites to be significantly enriched for motifs predicted to be binding sites for Zinc Finger (ZNF) transcription factors family members, as well as the transcription factor WT1 (**Figure 2D**). Cell-type-independent enhancers with H3K4me1 loss were enriched for the WT1 transcription factor motif, while cell-type-independent promoters with H3K4me1 loss were enriched for the EGR1 motif (**Supplementary Figure S5B-C**). KS1 sites that gained H3K4me1 peaks showed significant enrichment for KLF1 transcription factor binding site (**Figure 2D**).

Overall, these data show significant dysregulation of the H3K4me1 landscape in KS1 iPSCs, especially in the enhancer regions.

### The transcriptome is altered in KS1 iPSCs and is correlated with the H3K4me1 landscape

Altered H3K4me1 landscape is predicted to result in changes in the transcriptome. We, therefore, performed RNAseq on control and KS1 (c.16019G>A) iPSCs (**Supplementary Figure S7A**) and detected 909 significant DEGs (FDR<0.05, -1>Log2FC>1). Of these, 554 DEGs were downregulated while 355 were upregulated (**Figure 3A**). Gene Ontology (GO) overrepresentation analysis on the 909 DEGs detected biological categories related to organ morphogenesis and development, cell adhesion and actin and extracellular structure to be significantly overrepresented in DEGs (FDR<0.05) (**Figure 3B**). Notably, DEGs related to stem cell differentiation (*TBX2, TWIST1*), mesenchyme development (*GBX2, MEF2C*), negative regulation of locomotion (*ANGPT2, GREM1*), morphogenesis of an epithelium (*AJAP1, GDF7*) and regulation of neuron projection development (*POU3F2, NTRK2*) were upregulated while actin filament-based movement (*TNNC2*, *ACTA1*) and sensory perception of light stimulus (*TULP1*, *AIPL1*, *CRYBB1*) were downregulated (**Figure 3A**). GO categories for ear development (*TSHZ1*, *BMP5*), neuron projection (*GDF7*, *CNTN6*), sensory organ morphogenesis (*DIO3*, *NTRK2*), embryonic organ development (*PAX5*, *KDR*) and heart morphogenesis (*SNAI2*, *PITX2*) included both up and downregulated DEGs (**Figure 3A**).

**Figure 3.**
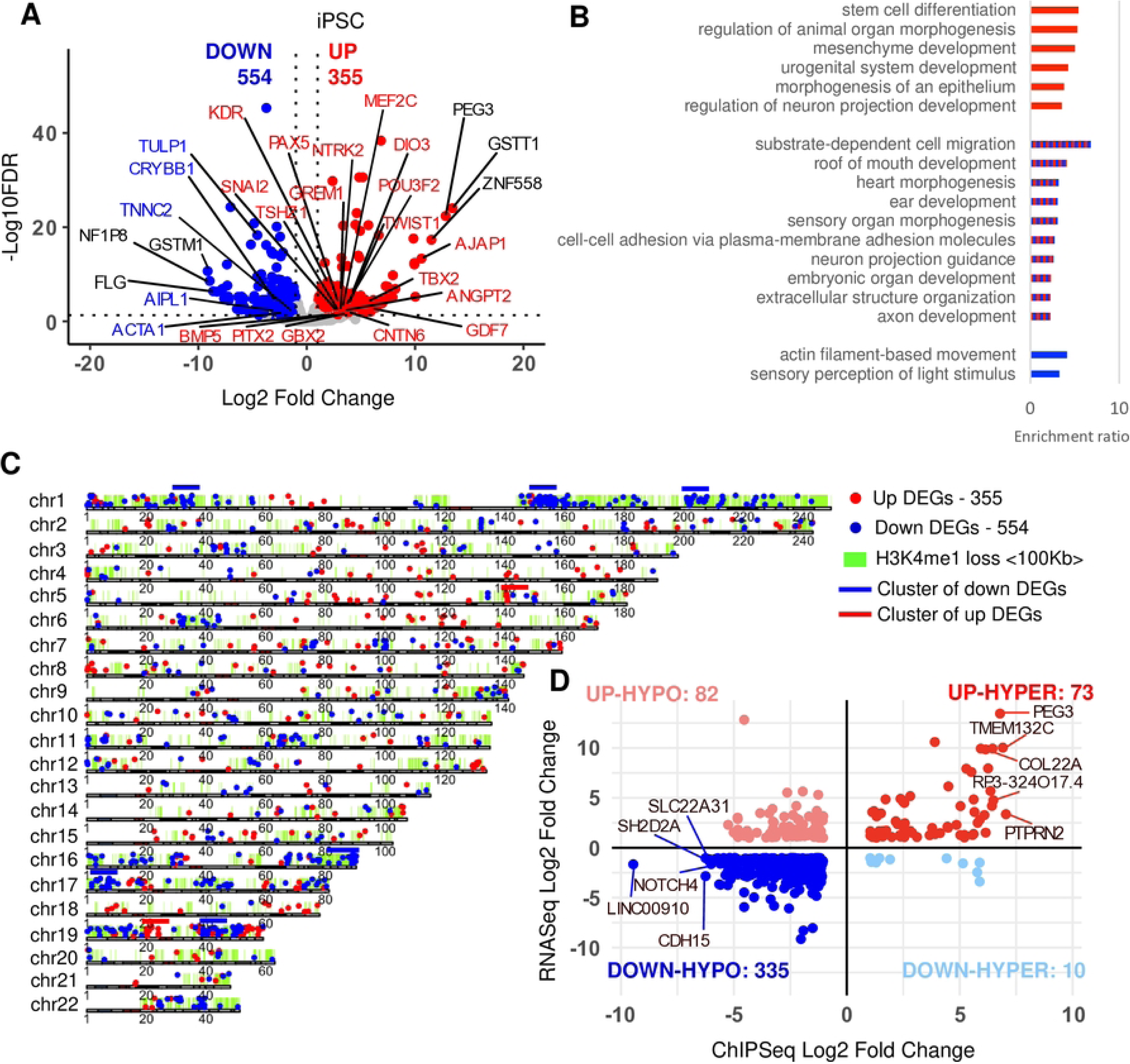
Correlation between changes in transcripts and H3K4me1 in KS1 iPSCs. **A)** Volcano plot showing down and up regulated genes on autosomal chromosomes in KS1 compared to controls (adjp<0.05, -1>log2FC>1). Genes highlighted are related to the GO categories in blue and red. Top 3 genes with highest or lowest log2FC are highlighted by black text. **B)** WebGestalt was performed on DEGs (adjp < 0.05; -1>log2FC>1) between KS1 iPSC and control iPSC. Gene ontology categories for biological processes are shown in the bar graphs (FDR<0.05). Red bars correspond to up DEGs, blue to down DEGs and red and blue to both. **C)** Karyoplot showing up (red) and down (blue) DEGs on each chromosome. Lime-green regions represent 100Kb up and downstream regions within H3K4me1 loss. Different grey colours represent the different G band staining and red represents centromere. Blue lines above chromosomes represent regions of enrichment of downregulated DEGs and red lines for upregulated DEGs (FDR < 0.05; hypergeometric test) **D)** Scatter plot showing change in expression and change in H3K4me1 within ±100kb of DEGs.

Next, we performed transcription regulator enrichment analysis on DEGs. This revealed SUZ12, a component of the Polycomb Repressive Complex 2 (PRC2), as the top regulatory factor among the upregulated DEGs (adjusted p-value < 0.05) (**Table 1**).

**Table 1.** Consensus Transcription factor (TF) analysis. CHEA and ENCODE consensus transcription factor (TF) analysis for upregulated (UP), downregulated (DOWN) and both up and downregulated (UP and DOWN) DEGs (adjp<0.05) at the three different differentiation stages and in the validation dataset.

| Stage | DEGs | Term | Overlap | P-value | Adj P-value |
| --- | --- | --- | --- | --- | --- |
| iPSC | UP | SUZ12 CHEA | 93/1684 | 8.92E-24 | 8.74E-22 |
|  |  | AR CHEA | 34/1095 | 0.001 | 0.055 |
|  |  | TP53 CHEA | 14/319 | 0.002 | 0.058 |
|  | DOWN | EZH2 ENCODE | 17/338 | 0.014 | 0.653 |
|  |  | REST CHEA | 49/1280 | 0.014 | 0.653 |
|  |  | REST ENCODE | 16/383 | 0.069 | 0.999 |
|  | UP and DOWN | SUZ12 CHEA | 144/1684 | 5.20E-14 | 5.26E-12 |
|  |  | SMAD4 CHEA | 38/584 | 0.017 | 0.708 |
|  |  | EZH2 ENCODE | 24/338 | 0.021 | 0.708 |
| Neural Progenitor | UP | SUZ12 CHEA | 31/1684 | 1.37E-04 | 0.012 |
|  |  | SALL4 CHEA | 7/355 | 0.045267 | 0.999996 |
|  |  | RAD21 ENCODE | 16/1265 | 0.118198 | 0.999996 |
|  | DOWN | GATA2 CHEA | 14/772 | 0.144067 | 0.999995 |
|  |  | ZEB1 ENCODE | 3/106 | 0.164940 | 0.999995 |
|  |  | STAT3 CHEA | 4/177 | 0.206557 | 0.999995 |
|  | UP and DOWN | SUZ12 CHEA | 48/1684 | 0.048317 | 0.999995 |
|  |  | STAT3 CHEA | 6/177 | 0.205444 | 0.999995 |
|  |  | KLF4 CHEA | 26/987 | 0.219578 | 0.999995 |
| Neuron | UP | SUZ12 CHEA | 15/1684 | 0.004000 | 0.245000 |
|  |  | EZH2 CHEA | 3/237 | 0.080143 | 0.999996 |
|  |  | KLF4 CHEA | 6/987 | 0.242292 | 0.999996 |
|  | DOWN | IRF8 CHEA | 1/121 | 0.171601 | 0.998295 |
|  |  | HDAC2 ENCODE | 1/175 | 0.238640 | 0.998295 |
|  |  | FOXA2 ENCODE | 1/316 | 0.389870 | 0.998295 |
|  | UP and DOWN | SUZ12 CHEA | 16/1684 | 0.033848 | 0.999996 |
|  |  | EZH2 CHEA | 3/237 | 0.159055 | 0.999996 |
|  |  | KLF4 CHEA | 7/987 | 0.348239 | 0.999996 |
| Neural Progenitor – validation datasets | UP | SUZ12 CHEA | 6/1684 | 0.00194 | 0.0329 |
|  |  | SUZ12 ENCODE | 1/105 | 0.0856 | 0.515 |
|  |  | REST CHEA | 3/1280 | 0.0909 | 0.515 |
|  | DOWN | GATA2 CHEA | 1/772 | 0.241 | 0.371 |
|  |  | NFE2L2 CHEA | 1/1022 | 0.307 | 0.371 |
|  |  | REST CHEA | 1/1280 | 0.371 | 0.371 |
|  | UP and DOWN | SUZ12 CHEA | 6/1684 | 0.0127 | 0.228 |
|  |  | REST CHEA | 4/1280 | 0.0638 | 0.547 |
|  |  | SUZ12 ENCODE | 1/105 | 0.119 | 0.547 |

Downregulation of several genes within a 110 Kb region on chromosome 19 was previously reported in KS1 iPSCs suggesting locus-specific targeting of KMT2D (16). We, therefore, asked if any clusters of DEGs were present in our dataset as well. We found significant enrichment of downregulated DEGs located on chromosome 1, 16, 17 and 19 (hypergeometric test, FDR< 0.0001) (**Figure 3C** and **Supplementary Figure S7C**). We also detected a region on chromosome 19, which overlaps with the previously reported downregulated cluster at 19q13.33 (16).

Next, we asked if there was correlation between the altered H3K4me1 signals and gene expression in our cell model. For this, we analysed expression of genes within ±100kb of differentially lost H3K4me1 regions. We found several regions of the genome with clusters of reduced H3K4me1 peaks and downregulated genes with significant enrichment of downregulated DEGs on chromosome 1, 16, 17 (hypergeometric test, FDR < 0.0001) (**Figure 3C** and **Supplementary Figure S7C**). (**Figure 3C**). We also found H3K4me1 and gene expression levels to be significantly correlated based on the ±100kb distance between gene TSS and H3K4me1 peak (p-value < 0.05; hypergeometric test) (**Figure 3D** and **Table 2**). Most of these genes correspond to areas near cell-type independent enhancer regions with loss of H3K4me1 (**Supplementary Figure S7D**).

**Table 2.** Hypergeometric analysis of gene-peak associations across different genomic distances. B) Association of DEGs and differential H3K4me1 peaks in iPSCs. **B)** Association of DEGs and differential H3K4me1 peaks in Neural Progenitors. **C)** Association of DEGs and differential H3K4me1 peaks in Neurons. Associations with p-value < 0.05 of downregulated DEGs and loss of H3K4me1 are in blue, and associations with p-value < 0.05 of upregulated DEGs and gain of H3K4me1 are in red.

A iPSC
| Peak set | Interval | iPSC_DGE_DOWN | p-value | iPSC_DGE_UP | p-value |
| --- | --- | --- | --- | --- | --- |
| iPSC_KSvsWT_FDR0.05_log2FC1_Gain | 5000 | 0 | 1 | 25 | 1E-12 |
| iPSC_KSvsWT_FDR0.05_log2FC1_Gain | 25000 | 4 | 0.987119325 | 41 | 1E-12 |
| iPSC_KSvsWT_FDR0.05_log2FC1_Gain | 50000 | 8 | 0.990802013 | 50 | 1.40546E-11 |
| iPSC_KSvsWT_FDR0.05_log2FC1_Gain | 100000 | 16 | 0.995582584 | 63 | 3.66108E-11 |
| iPSC_KSvsWT_FDR0.05_log2FC1_Gain | 150000 | 26 | 0.988047687 | 76 | 5.39185E-11 |
| iPSC_KSvsWT_FDR0.05_log2FC1_Gain | 200000 | 29 | 0.999412021 | 84 | 1E-12 |
| iPSC_KSvsWT_FDR0.05_log2FC1_Loss | 5000 | 110 | 2.74261E-11 | 4 | 0.999980277 |
| iPSC_KSvsWT_FDR0.05_log2FC1_Loss | 25000 | 214 | 1.78334E-11 | 21 | 1 |
| iPSC_KSvsWT_FDR0.05_log2FC1_Loss | 50000 | 256 | 1E-12 | 39 | 1 |
| iPSC_KSvsWT_FDR0.05_log2FC1_Loss | 100000 | 308 | 1E-12 | 62 | 1 |
| iPSC_KSvsWT_FDR0.05_log2FC1_Loss | 150000 | 338 | 2.02831E-11 | 84 | 1 |
| iPSC_KSvsWT_FDR0.05_log2FC1_Loss | 200000 | 351 | 2.69212E-09 | 95 | 1 |

B Neural progenitor
| Peak set | Interval | Prog_DGE_DOWN | p-value | Prog_DGE_UP | p-value |
| --- | --- | --- | --- | --- | --- |
| Prog_KSvsWT_FDR0.05_log2FC1_Gain | 5000 | 14 | 0.200926 | 37 | 1E-12 |
| Prog_KSvsWT_FDR0.05_log2FC1_Gain | 25000 | 35 | 0.631637 | 58 | 4.52E-10 |
| Prog_KSvsWT_FDR0.05_log2FC1_Gain | 50000 | 52 | 0.934551 | 74 | 1.98E-07 |
| Prog_KSvsWT_FDR0.05_log2FC1_Gain | 100000 | 74 | 0.999698 | 88 | 0.002872 |
| Prog_KSvsWT_FDR0.05_log2FC1_Gain | 150000 | 95 | 0.999985 | 99 | 0.073142 |
| Prog_KSvsWT_FDR0.05_log2FC1_Gain | 200000 | 113 | 0.999996 | 107 | 0.304067 |
| Prog_KSvsWT_FDR0.05_log2FC-1_Loss | 5000 | 19 | 0.00581 | 2 | 0.993048 |
| Prog_KSvsWT_FDR0.05_log2FC-1_Loss | 25000 | 62 | 0.013411 | 21 | 0.993861 |
| Prog_KSvsWT_FDR0.05_log2FC-1_Loss | 50000 | 92 | 0.139876 | 36 | 0.999894 |
| Prog_KSvsWT_FDR0.05_log2FC-1_Loss | 100000 | 126 | 0.83926 | 55 | 1 |
| Prog_KSvsWT_FDR0.05_log2FC-1_Loss | 150000 | 137 | 0.999887 | 71 | 1 |
| Prog_KSvsWT_FDR0.05_log2FC-1_Loss | 200000 | 151 | 0.999999 | 81 | 1 |

C Neuron
| Peak set | Interval | Neur_DGE_DOWN | p-value | Neur_DGE_UP | p-value |
| --- | --- | --- | --- | --- | --- |
| Neur_KSvsWT_FDR0.05_log2FC1_Gain | 5000 | 1 | 0.431159 | 8 | 0.000151 |
| Neur_KSvsWT_FDR0.05_log2FC1_Gain | 25000 | 1 | 0.894369 | 17 | 7.05E-05 |
| Neur_KSvsWT_FDR0.05_log2FC1_Gain | 50000 | 3 | 0.768472 | 22 | 0.000684 |
| Neur_KSvsWT_FDR0.05_log2FC1_Gain | 100000 | 3 | 0.977701 | 26 | 0.037156 |
| Neur_KSvsWT_FDR0.05_log2FC1_Gain | 150000 | 5 | 0.977017 | 30 | 0.178229 |
| Neur_KSvsWT_FDR0.05_log2FC1_Gain | 200000 | 5 | 0.996821 | 37 | 0.117381 |
| Neur_KSvsWT_FDR0.05_log2FC1_Loss | 5000 | 3 | 0.598051 | 5 | 0.928874 |
| Neur_KSvsWT_FDR0.05_log2FC1_Loss | 25000 | 6 | 0.877893 | 12 | 0.998469 |
| Neur_KSvsWT_FDR0.05_log2FC1_Loss | 50000 | 10 | 0.867032 | 18 | 0.99995 |
| Neur_KSvsWT_FDR0.05_log2FC1_Loss | 100000 | 14 | 0.949907 | 27 | 1 |
| Neur_KSvsWT_FDR0.05_log2FC1_Loss | 150000 | 14 | 0.998383 | 34 | 1 |
| Neur_KSvsWT_FDR0.05_log2FC1_Loss | 200000 | 15 | 0.999797 | 40 | 1 |

Overall, these data show that at the stem cell stage there are significant changes in expression of genes related to the KS1 phenotype, which are over-represented by regions of DNA associated with SUZ12 recruitment.

### The neuronal progenitor and neural epigenomes are altered in KS1

Having shown the epigenomic changes at the stem cells stage, we asked if similar changes could also be present in differentiating cells. Learning difficulties are seen in 95% individuals and epilepsy is present in 25% individuals affected by KS1(9, 18). We, therefore, differentiated the KS1 (c.16019G>A) and control iPSCs into neuroepithelium after 10 days of SMAD signalling inhibition (19) giving rise to neural rosettes (**Supplementary Figure S8A**) and later into early cortical neurons (Day 30) (**Figure 4A** and **Supplementary Figure S9A**). After the formation of the rosettes, cells were enriched for progenitors by FACs for a CD44^-^CD184^+^CD24^+^ population at Day 18 and for neuronal cells by sorting for a CD44^-^CD184^-^CD24^+^ population at Day 30 (**Figure 4A**). Both control and KS1 cells expressed key markers of neuronal progenitors such as *PAX6*, *FOXG1, SOX2, NES* and *OTX2*, and of neuronal cells such as *TBR1, MAP2* and *GRIA2* (**Figure 4B**).

**Figure 4.**
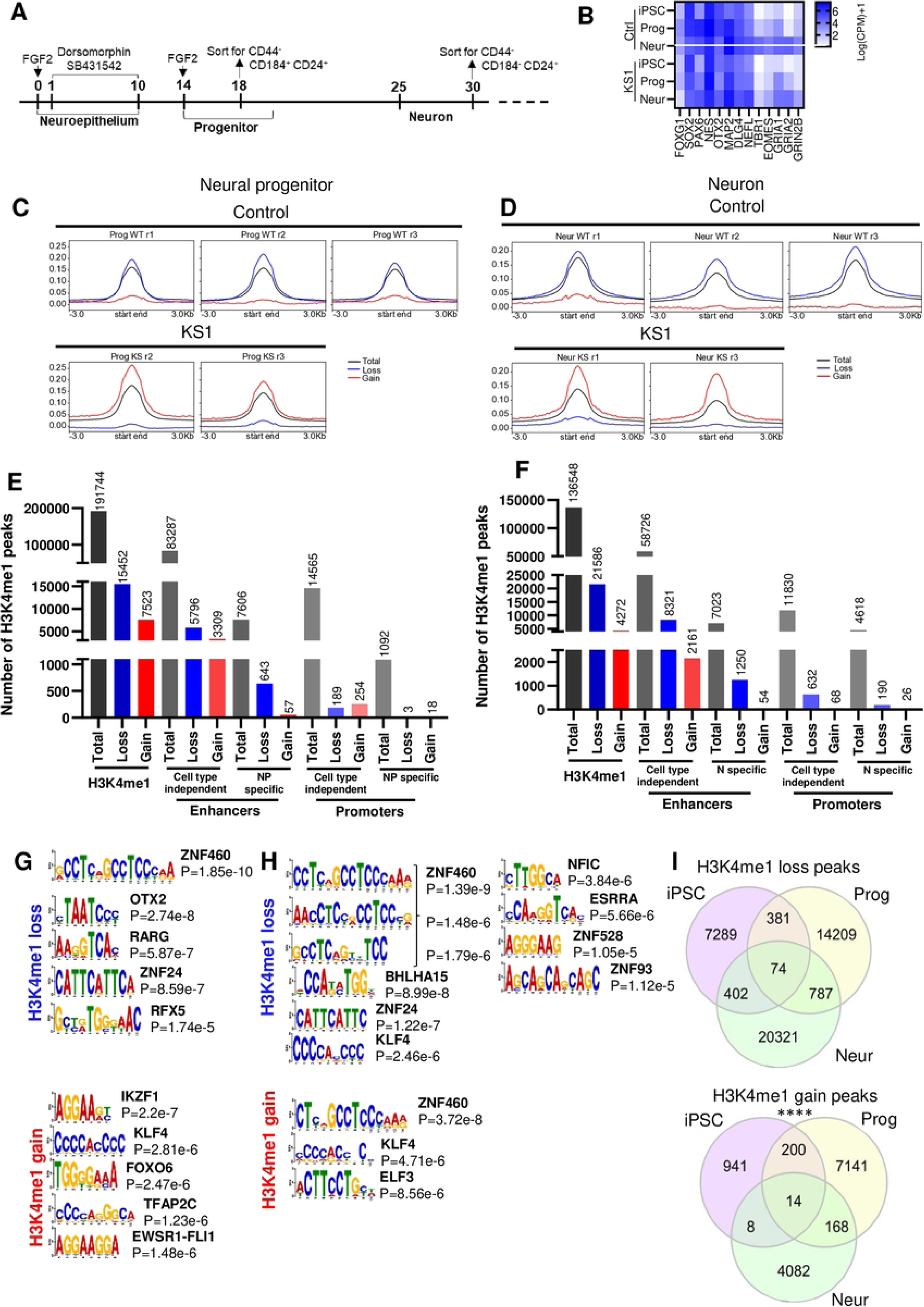
Loss of H3K4me1 in KS1 neural progenitors and neurons. **A)** Schematic representation of the neuronal differentiation protocol **B)** RNAseq analysis showing transcript level for key neural progenitor markers in control and KS1 samples at stem cell, progenitor and neuronal stages. **C-D)** Line graph of H3K4me1 peaks (black), H3K4me1 loss regions (blue), and H3K4me1 gain regions (red) for neuronal progenitors and neurons. **E-F)** Bar graph showing number of H3K4me1 peaks in relation to enhancer and promoter regions for total, loss and gain in mono-methylation. **G-H)** Top TF motifs predicted in differential H3K4me1 peaks (loss and gain of H3K4me1 in binding regions) between KS1 and control. **I)** Intersection analysis showing H3K4me1 peaks in common between iPSC, Progenitor and Neurons (hypergeometric test; ****p<0.0001) between KS1 (c.16019G>A) and control.

We performed H3K4me1 ChIPseq on sorted neural progenitors and neuronal cells. In neural progenitors, we detected 22,975 differentially methylated sites (FDR<0.05; -1>Log2FC>1). Of these, 15,452 (67.2%) sites showed loss and 7,523 showed gain in H3K4me1 in KS1 cells (**Figure 4C-4E** and **Supplementary Figure S8**). In neurons, we detected 25,858 differentially methylated sites (FDR<0.05; -1>Log2FC>1). Of these, 21,586 sites showed loss in H3K4me1 and 4,272 showed gain in H3K4me1 (**Figure 4D-4F** and **Supplementary Figure S9**). Of the H3K4me1 peak site losses, 5,796 (37.5%) in neural progenitor and 8,321 (38.5%) in neurons correspond to cell-type independent enhancer regions (**Figure 4E-F** and **Supplementary Figure S6B-C**).

Motif analysis on neuronal progenitors revealed that the sites where H3K4me1 was lost in KS1 cells were predicted to target for ZNF family of transcription factors as well as OTX, RARG and RFX5 while sites where H3K4me1 was gained were predicted to be targets for IKZF1, KLF4, FOXO6, TFAP2C and EWSR1-FLI1 (**Figure 4G**). ZNF460, PATZ1, PITX1 and ZKSCAN1 were found to be associated with cell-type independent enhancers with H3K4me1 loss while ELF1, TFAP2C and KLF1 with cell-type independent enhancers with H3K4me1 gain (**Supplementary Figure S8D**). In neuronal cells, loss of H3K4me1 sites were predicted to be bound by ZNF transcription factor family members, BHLHA15, KLF4, NFIC and ESRRA (**Figure 4H**). ZNF transcription factor family members, NEUROD1, ARGFX, NFIB and NFIC motifs were found to be associated with cell-type independent enhancers with H3K4me1 loss (ZNF was also found in cell-type independent promoters with H3K4me1 loss) (**Supplementary Figure S9D-E**).

Next, we compared the significantly differentially methylated peaks across the three stages of neuronal differentiation. We identified 88 H3K4me1 regions (74 losses and 14 gains) that were affected across all stages in the KS1 lines (**Figure 4I**). Three hundred H3K4me1 loss peaks were maintained between iPSC and neural progenitor stage and more than 700 between neural progenitor and neuronal cells stage (hypergeometric test; p-value<0.0001) (**Figure 4I**).

Overall, in our KS1 model we witness increasing dysregulation of the H3K4me1 landscape with advancing differentiation from iPSCs to neurons.

### The transcriptome is altered in KS1 neuronal progenitors and neurons

Finally, we investigated the transcriptome of KS1 neuronal progenitors and neurons by RNAseq (**Supplementary Figure S7B**). In neuronal progenitors, we detected 447 DEGs (FDR<0.05, - 1<LOGFC>1). Of these, 264 DEGs were downregulated while 183 DEGs were upregulated in KS1 neuronal progenitors (**Figure 5A**). GO analysis on the 447 DEGs revealed biological categories related to organ development, cell fate commitment and extracellular structure (FDR<0.05) (**Figure 5C**). Notably, DEGs related to neuron projection guidance (*FEZ1*, *VANGL2*) and cilium organization (*CDK10*, *DNALI1*) were downregulated whereas morphogenesis of epithelium (*AJAP1, IRX2*) and embryonic organ development (*TBX1, EN2*) were upregulated. GO categories for urogenital system development (*PAX8*, *AMH*), negative regulation of nervous system development (*INPP5F*, *SOX8*), and synapse organisation (*SLC7A11*, *NTNG2*) were contributed by both up- and downregulated DEGs. In neuronal cells, we detected a lower number of DEGs with only 116 DEGs (FDR<0.05, -1>Log2FC>1) and 147 DEGs (FDR<0.1, -1>Log2FC>1) (**Figure 5B**). Of these, 31 (44; FDR<0.1) DEGs were downregulated while 85 (103; FDR<0.1) DEGs were upregulated (**Figure 5B**). GO analysis on the 147 DEGs (FDR<0.1) showed biological categories related to related to nerve development (*TBX1*, *NGFR, EMX1*) contributed by both up and down DEGs while GO categories related to pattern specification process (*HOXB2*, *HOXB3*, *BARX1*), autonomous nervous system development (*SOX10*, *TBX1*) were upregulated (**Figure 5D**).

**Figure 5.**
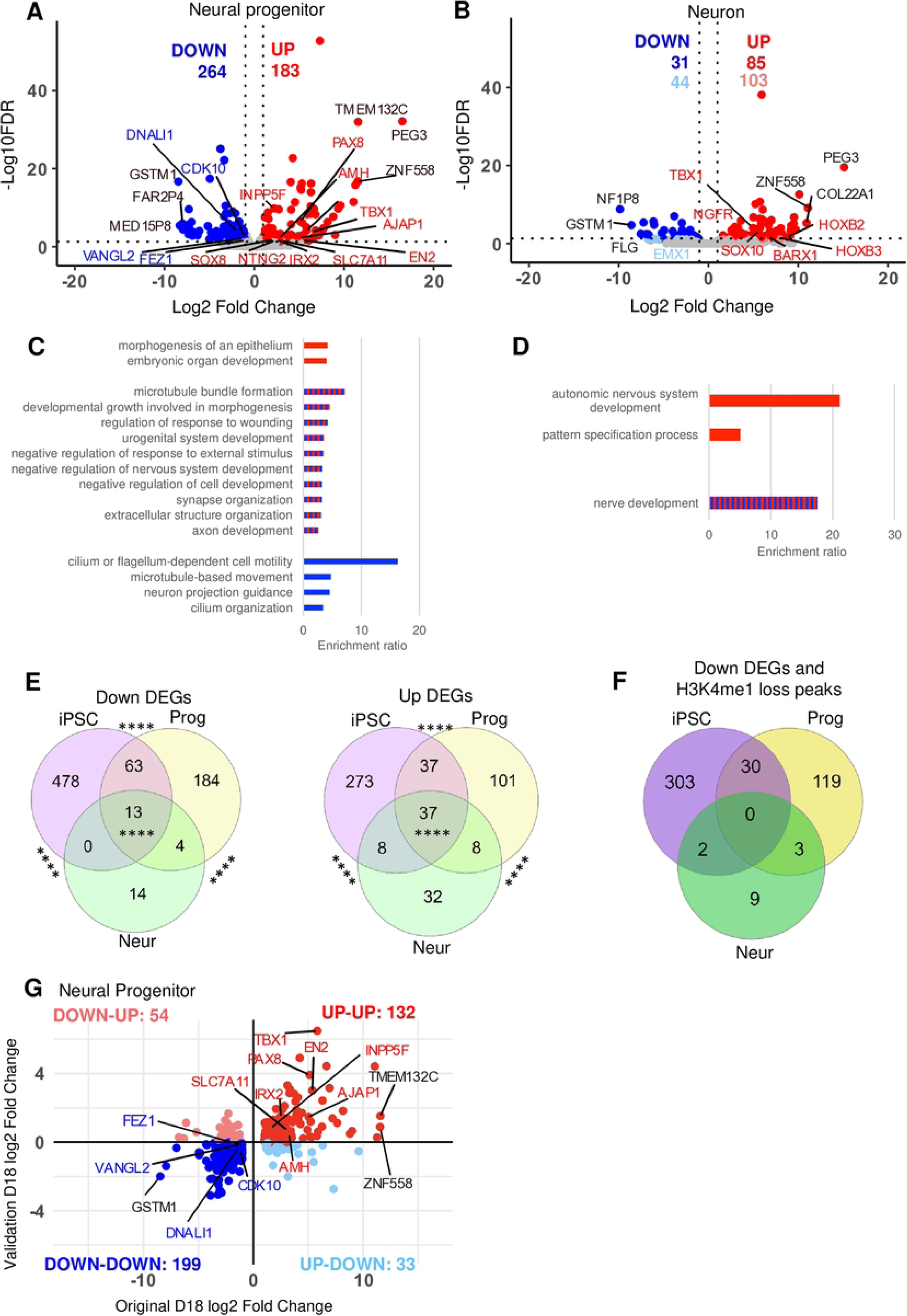
Changes in the transcriptome in KS1 neural progenitors and neurons and comparison of transcriptomic and epigenomic changes during neuronal differentiation. A-B) Volcano plot showing down and up regulated genes on autosomal chromosomes in KS1 compared to controls (adjp<0.05, -1>log2FC>1 in red/blue; adjp<0.1, -1>log2FC>1 in light-red/light-blue). Top 3 genes with highest or lowest log2FC are highlighted in black. **C-D)** WebGestalt was performed on DEGs (adjp<0.1, -1>log2FC>1) between KS1 and control in neuronal progenitor and neuronal cells. Gene ontology categories for biological processes are shown in the bar graphs (FDR<0.1) Red bars correspond to upregulated (up) DEGs, blue to downregulated (down) DEGs and red and blue to both. **E)** Intersection analysis showing DEGs in common between iPSC, progenitor and neurons between KS1 (c.16019G>A) and control (exact test; ****p<0.0001). **F)** Intersection analysis showing Down DEGs and H3K4me1 loss peaks in common between iPSC, progenitor and neurons between KS1 (c.16019G>A) and control. **G)** Comparison of Log2 Fold Changes of DEGs from the original dataset (DS1) and validation dataset (DS3) at progenitor stage (Day18). Genes within Gene Ontology categories are highlighted in blue and red. Top 3 genes with highest or lowest log2FC are highlighted in black.

Transcription factor enrichment analysis revealed SUZ12 as top regulatory factor for upregulated DEGs in neural progenitors (adjusted p-value < 0.05) (**Table 1**). SUZ12 was also the top most hit for neurons, although not statistically significant. Additionally, we found significant enrichment of downregulated DEGs located on chromosome 1, 16, 17 in neural progenitors and in 1 and 5 in neuronal cells (hypergeometric test, p-value < 0.0001) (**Supplementary Figure S10-11**).

Next, to analyse which changes in the transcriptome were in common at all three stages of neuronal differentiation, we compared DEGs between the three differentiation stages. Interestingly, we observed 50 significant DEGs (13 down- and 37 upregulated) in common across all three differentiation stages (**Figure 5E**). Intersection analysis showed most genes were significantly dysregulated between the iPSC and neural progenitor stage (exact test; p<0.0001).

To identify possible direct associations between H3K4me1 loss and downregulated genes across multiple stages of differentiation in our model, we extracted genes, which are downregulated and have transcription start sites within ±100kb of H3K4me1 loss. We identified 30 such genes in common between iPSC and neural progenitor stages, of which 4 (*CNTN4*, *NOVA2*, *GAS8* and *TRIM46*) are involved into axogenesis, neuronal projection guidance and microtubule formation (**Figure 5F**).

Overall, in our KS1 model we witness a decrease in the number of DEGs with advancing neuronal differentiation.

### The altered KS1 transcriptome is consistent

Most of our neuronal differentiation work was based on a single missense mutant and we observed the biggest differences in the Day 0 and Day 18 transcriptomes (**Figure 5E**). We, therefore, decided to validate our results via RNAseq at Day 18 in neural progenitors from our original control and mutant lines, and in two additional controls and patient-derived iPSCs with distinct mutations (c.5527dupA, c.16180G>T). We found several up- and downregulated DEGs to follow the same gene expression trend (adjusted p-value<0.05; -1<LOG2FCϬ1) (**Figure 5G**). Furthermore, 331/447 DEGs from our original Day 18 DEGs dataset (**Figure 5A**) have the same gene expression trend in KS1 samples as in the validation Day 18 DEGs dataset (Spearman correlation; rho = 0.61; p-value < 0.0001) (**Figure 5G**). Additionally, transcription factor enrichment analysis performed on the Day 18 validation dataset confirmed SUZ12 as a key transcriptional regulator for upregulated DEGs reinforcing its main role in gene regulation in KS1 (**Table 1**).

Overall, these results validate our findings in additional cell lines harbouring distinct *KMT2D* mutations.

## DISCUSSION

Although the underlying disease mechanism of KS1 has been examined previously(7, 15, 16), several questions about the pathological mechanism of the neurological issues in the disorder remain unanswered. Here, we studied patient-derived iPSC models to interrogate the basis of neurological phenotypes of KS1.

Firstly, haploinsufficiency has been the proposed mechanism for nonsense or frameshift variants in KS1, but it has not been previously proven. We observed ALT:REF read ratios to be significantly lower in iPSCs with nonsense or frameshift variants in comparison with the iPSCs with missense variants (**Figure 1B**). This observation suggests partial nonsense mediated mRNA decay of the mutant transcripts with premature termination codon. Interestingly, the absolute level of REF allele counts was higher in iPSCs with nonsense or frameshift variants than the ones with missense variants, which is likely explained by mRNA decay triggered transcriptional adaptation(20). The lower levels of reference transcripts in mutant cells with nonsense or frameshift variants in comparison with the controls is also reflected in the protein levels (**Figure 1C**). The KMT2D protein levels were also significantly lower in iPSCs with the missense variant, even though the overall transcript levels in these lines were not significantly different from controls. This indicates possibly reduced stability of the protein with the amino acid substitution. The p.(Arg5340GIn) variant is located in the FYR-N motif which may alter the total charge of the domain and/or the protein stability(21). It is possible that other KS1 causing missense variants could cause loss of function through other mechanisms such as altered protein-protein or protein-chromatin interactions or loss of enzyme activity or mislocalization. In contrast with a previous report of decrease in H3K4me3 levels in a KS1 mouse model(7), but in line with the known role of KMT2D as a H3K4 mono-methyltransferase(1, 3), we found a significant decrease in overall cellular H3K4me1 levels at the iPSC stage (**Figure 1D**). This is also in line with the finding of analysis of 33 KS1 and 36 healthy controls’ enhancer signatures related to H3K4me1 and H3K4me2 in peripheral blood mononuclear cells of patients with KS1(14).

Our experiments also show dysregulation of H3K4me1 and transcriptome landscapes in KS1 iPSCs, neuronal progenitors and neurons (**Figures 2**-**5**). The vast majority of differentially methylated H3K4me1 peaks were losses, which is in line with the known role of KMT2D as a H3K4 methyltransferase(1, 2). Importantly, we find significant correlation between the H3K4me1 and transcriptome dysregulation. This suggests that several of the downregulated DEGs are likely to be direct consequences of loss of H3K4me1 rather than secondary to network disruption. We suspect that these correlations in KS1 may be an underestimate as we only considered regions 100Kb up and downstream of DEGs as well as H3K4me1 peaks with ±1 fold change for our analysis, which ignores potential long-range enhancer activity and potential *trans*-acting effects of H3K4me1 deposition. Furthermore, we observed ‘large blocks of dysregulation’ rather than it being evenly spread throughout the genome suggesting that certain regions of the genome are more susceptible to loss of KMT2D. Interestingly, with progress in differentiation, we observed an increase in loss of H3K4me1 peaks but decrease in downregulated DEGs. This could reflect higher dependency of H3K4me1 regulation on KMT2D at the pluripotent stage. Network level adaption could be an alternative explanation.

The analysis of transcription factor binding sites in our ChIPseq dataset revealed several transcription factors that may be important in KS1 pathology **(Figures 2 and 4)**. KLF(22) and ZNF (23) family transcription factors play crucial roles in cellular processes such as cell growth, differentiation, apoptosis, and response to environmental stimuli. Wilms Tumour 1 (WT1) expression is required in early kidney development and pathogenic mutations in *WT1* gene are associated with genitourinary phenotypes(24) Estrogen-related receptors (ESRR) regulate genes involved in energy homeostasis, including fat and glucose metabolism and mitochondrial biogenesis(25). Regulatory factor X-5 (RFX5) is involved in regulation of major histocompatibility class II and *RFX5* mutations result in immunodeficiency(26). IKAROS family zinc finger 1 (IKZF1) is involved in lymphocyte development and pathogenic variants in IKZF1 are associated with immunodeficiency(27). OTX2 is critical for early embryonic development, particularly in the development of the brain and sensory organs(28). The effects of retinoic acid on gene expression is mediated by RARG, influencing various biological processes, including development, differentiation, and homeostasis(29). FOXO6 is involved in regulation of Hippo signalling during craniofacial development, with FoxO6^-/-^ have facial overgrowth(27). ETS variant (ETV) transcription factors are required for hippocampal dendrite development(30), while TFAP2C is vital for early development, specifically in morphogenesis(31). Additionally, ETS protein can regulate immune responses and the development of immune-related cells(32).

In our RNAseq datasets, we detected several DEGs important in lineage commitment and neuronal differentiation (**Figures 3 and 5**). In particular, *PAX8,* important in tissue and organ formation particularly in kidney, thyroid gland and brain(33), is upregulated in KS1 neural progenitor cells. *PAX5,* involved in early neural tube development and regionalization(34), is also upregulated in KS1 iPSC and neural progenitor cells. Its dysregulation in KS1 cells might affect the correct formation of the neural tube, contributing to the KS1 intellectual disability phenotype. We also found SUZ12 to be a master regulator of many of the upregulated DEGs in our datasets (**Table 1**). SUZ12 is a component of PRC2 required for the H3K27 tri-methylation mark(35). Further studies will be required to clarify the complex relationships between the chromatin and transcriptional landscape in KS1.

In conclusion, these results establish heterozygous truncating or missense *KMT2D* variants cause KS1 through a loss of function mechanism and result in significant dysregulation of H3K4me1 and the transcriptome landscape across different stages of neuronal differentiation. The H3K4me1 and transcriptome landscapes of KS1 are correlated and their dysregulation seem to occur in discreet blocks across the genome. With advancing differentiation from iPSCs to neurons, the dysregulation of H3K4me1 landscape increase but the number of DEGs decrease.

## MATERIAL AND METHODS

### Generation, validation and culture of induced pluripotent stem cells

Skin biopsies and blood samples were obtained from individuals using standard procedures. Fibroblast cells were cultured in DMEM, 10% foetal bovine serum, 1% L-glutamine at 37°C in a humidified 5% CO_2_ incubator. Fibroblast cells or peripheral blood mononuclear cells (for Ctrl 2) were reprogrammed into iPSC by using Sendai viral vectors (CytoTuneTM −iPS 2.0 Sendai Reprogramming Kit-ThermoFisher). Control iPSC (wigw2 - Ctrl 1) and KS1 iPSCs (ierp4 - iPSC 3, aask4 - iPSC 1, oadp4 - iPSC 2) were generated by HipSci (https://www.hipsci.org/#/cohorts/kabuki-syndrome) while control iPSCs (JF191b(36) - Ctrl 2, SW171a(37) - Ctrl 3) were generated in house under local ethics (REC 11/H1003/3; IRAS ID 64321 or REC 18/EE0250 IRAS246779). All were grown on Vitronectin N-terminal fragmented (Life Technologies) coated dishes, using Essential 8 medium (StemCELL Technologies). Cells were routinely passaged when 80% confluent using 0.1% EDTA-PBS without Ca^2+^ and Mg. All cell lines were cultured at 37°C in a humidified 5% CO_2_ incubator.

### Neuronal differentiation

HiPSCs were differentiated to cortical neuron using a protocol adapted from Shi et al 2012(19). At the start of the protocol, on Day 0, cells were plated on Matrigel (Corning) and cultured using E8 medium (StemCell Technologies) supplemented with FGF2 (10ng/ml). On Day 1, neuronal differentiation was undertaken using neuronal induction medium supplemented with SMAD signalling inhibitors, Noggin (500ng/ml, Tocris) or Dorsomorphin (Abcam) and SB43154 (10uM, Tocris). On Day 10, cells were passaged and plated on a poly-l-ornithine (Sigma) and murine laminin L2020 (20μg/ml, Sigma) substrate for neuronal maintenance and maturation. On Day 12, small-elongated cells generate rosette structures resembling early neural tubes which were propagated in culture using FGF2 (20ng/ml) for about 3 days. On Day 18, cells were passaged using Dispase (StemCell Technologies), plated on a poly-l-ornithine and laminin substrate for neuronal maturation and cultured up to Day 30.

### Fluorescence-activated cell sorting (FACS)

Single cell suspensions from neural progenitor (Day 18) and neuronal (Day 30) culture were stained and sorted with an Aria Fusion flow cytometer (BD Biosciences). Cells were blocked in PBS-5%FBS for 30 minutes and incubated with cell surface antibodies, CD44-PE (561858), CD24-FITC (560992) and CD184-APC (560936) for 30 minutes at 4°C. After staining, cells were washed in PBS-5%FBS and filtered through a 0.22 µm filter before sorting and acquisition. All antibodies used for staining were purchased from BD Bioscience. Cells used for RNAseq and ChIPseq were sorted for CD44^-^ CD24^+^ and CD184^+^ for the progenitor stage at Day 18 and for CD44^-^ CD24^+^ and CD184^-^ (BD Bioscience) for neurons at Day 30.

### Immunofluorescence staining

Cells grown in 12-well plates were fixed with 4% paraformaldehyde for 15 minutes at room temperature. Samples were permeabilised with 0.5% Triton X-100 in PBS for 10 minutes at room temperature. Then blocking solution (1% BSA in PBS) was applied for 10 minutes at room temperature. Blocking solution was removed and an antibody recognising one of NANOG, OCT4, or SOX2 (4903S, 2890S, 3579S, Life Technologies) was applied overnight at 4°C. After washing three times with PBS, cells were incubated with secondary antibodies of Alexa FluorTM 488 goat anti-rabbit (A11008, Invitrogen) for 45 minutes at room temperature in the dark. After washing three times with PBS, samples were stained with DAPI (4083S, Cell signalling technology) for 5 minutes. Finally, samples were washed with PBS. Pictures were taken using an epifluorescence microscope (Olympus U-LH100HG) and cellSens software. Images are representative of three independent experiments.

### Western blot

Nuclear protein samples were isolated from iPSCs using NE-NER^TM^ Nuclear and Cytoplasmic Extraction Reagents (ThermoFisher Scientific, Cat#: XJ355181). Then protein samples were quantified using BCA™ protein assay kit-reducing agent compatible (ThermoFisher Scientific, Cat#: 23250). Electrophoresis was carried out using 10% Bis-Tris Gels and NuPAGE MES SDS Running Buffers (Invitrogen) or 3∼8% Tris-Acetate Gels (Invitrogen) and Tris-Acetate SDS Running Buffer (Life Technologies) for KMT2D. Twenty to forty µg of nuclear protein extracts were loaded into the polyacrylamide gel and electrophoresed for 45-60 minutes at 120V. Gels were blotted onto nitrocellulose membranes using Mini iBlot Gel Transfer Stacks Nitrocellulose (Invitrogen) for H3K4me1, me2 and me3, or wet transferred at 200mA over night at 4°C using transfer buffer (pH 8.3) contained glycine, tris base, sodium dodecyl sulfate, methanol and water. After blocking non-specific binding, the membranes were incubated with specific anti-H3K4me1 (ab8895, Abcam), anti-H3K4me2 (C15410035, Diagenode), anti-H3K4me3 (ab8580, Abcam), anti-KMT2D (ab213721, Abcam), and anti-HSP90 (4874S, Cell Signalling) overnight at 4°C. Then, the membrane was incubated with a secondary fluorescent labelled goat anti-rabbit antibody (IRDye 800CW LiCor) and the signal was developed using an Odyssey M imaging machine. Images were analysed using ImageJ by measuring the integrated density of the bands. Three experimental replicates were performed.

### Gene expression analysis

Cells were collected by centrifugation and total RNAs were extracted using RNeasy Mini kit (Qiagen) according to the manufacturer’s protocol. RNA concentration was measured using a NanoDrop 2000 spectrophotometer (Thermo Scientific). One μg of RNA was reverse transcribed with random hexamers primer (Promega) to generate cDNA using the M-MLV Reverse Transcriptase kit (Promega), according to manufacturer’s protocol. Quantitative real-time PCR (qRT-PCR) reactions were performed in triplicate on a Bio-Rad CFX394 Real Time system (Bio-Rad) using Power SYBR Green PCR Master mix (Applied Biosystems). For each sample, 2 µl cDNA (2 ng/µl) was incubated in a final volume of 10 µl with 5 µl of Power SYBR Green PCR Master mix (Applied Biosystems) and 0.5 µl of both the forward and reverse target-specific primer (10 µM, Sigma). The expression of each target gene was evaluated using a relative quantification approach (2^-ΔCT^ method) with GAPDH as the internal reference for human genes.

### RNA sequencing and analysis

RNAs were extracted using RNeasy Mini kit (Qiagen) according to the manufacturer’s protocol. Library preparation was performed by the Genomic technologies core facility at the University of Manchester. Samples were sequenced using NextSeq500 (Dataset 1 – DS1, Dataset 3 – DS3) or Illumina HiSeq 2000 sequencer (Dataset 2 from HipSci consortium – DS2). Unmapped paired reads were output in BCL format and converted to FASTQ format using bcl2fastq v2.17.1.14. Prior to the alignment, RNAseq reads were trimmed to remove any adapter or poor-quality reads using Trimmomatic v0.36 with options: “ILLUMINACLIP:Truseq3-PE-2_Nextera-PE.fa:2:30:10 SLIDINGWINDOW:4:20 MINLEN:35”(38). The filtered reads were mapped to the human reference genome (hg19/GRCh37) with comprehensive Gencode v19 genome annotation using STAR v2.5.3a with default options(39). Duplicate reads were marked using Samtools 1.9(40) and sorted by name. Reads were attributed to genes using FeatureCounts tool in Subread v2.0.0(41) in paired-end mode. Prior to the differential gene expression analysis, genes which have less than 10 reads in >3 samples were filtered out, as well as genes on sex chromosomes. Differential gene expression analysis was performed using DESeq2 v1.40.2 using Wald test with Benjamini Hochberg p value adjustment (p adjusted<0.05). EdgeR was used to calculate counts per million. GO over-representation analysis was performed using WebGestalt tool (2019 version). EnrichR(42, 43) was used for transcription regulator enrichment analysis. PEGS(44) tool was used to calculate enrichment of differentially expressed genes adjacent to differential ChIP-Seq H3K4me1 peaks. Statistical analysis of overlap of differentially expressed genes across timepoints was performed using SuperExactTest package within R v4.3.1. To assess clustering of DEGs in a statistical manner, we performed hypergeometric test for over-enrichment of DEGs in 10Mb genomic bins across autosomes (excluding patches, haplotypes and undefined contigs).

For validation heatmap generation and gene-based Z-score transformation of RNASeq count data pheatmap package within R v4.3.1 was used. Separate DGE analysis was performed for validation dataset 3 (-1<LOG2 FC>1, adjusted p-value <0.05). Then DEGs from Dataset 1 were intersected with DEGs from validation Dataset 2 or 3 and plot them in a heatmap representation.

### Chromatin Immunoprecipitation Sequencing and Analysis

ChIPseq has been performed using the True MicroChIP kit from Diagenode following the manufacturer’s protocol. Briefly, cells were collected and DNA-protein was cross-linked using 36.5% formaldehyde. Then cells were lysed and chromatin was sheared by sonication (Bioruptor NextGen). Sheared chromatin was incubated with H3K4me1 Antibody (5326, Cell signalling) overnight at 4°C. Library preparation was performed by the Genomic technologies core facility at the University of Manchester.

Unmapped paired-end sequences from an Illumina HiSeq4000 sequencer were output in BCL format and converted to FASTQ format using bcl2fastq v2.20.0.422, during which adapter sequences were removed. Unmapped paired-reads of 76 bp were interrogated using a quality control pipeline consisting of FastQC v0.11.3 (http://www.bioinformatics.babraham.ac.uk/projects/fastqc/) and FastQ Screen v0.9.2(45). Prior to the alignment, reads were trimmed to remove any adapter or poor-quality sequence using Trimmomatic v0.36(38), and truncated at a sliding 4bp window, starting 5’, with a mean quality <Q20, then removed if the final length was less than 35bp. The filtered paired-reads were mapped to the human reference sequence analysis set (hg19/GRCh37) from the UCSC browser, using Bowtie2 v2.3.0(46). Mapped reads were then filtered using Samtools v0.1.19 to retain only high-confidence concordant pairs (-f 2 -q30), followed by the removal of reads mapping to the mitochondrial genome and to unassembled parts of the reference genome. Peak calling was performed using MACS2 v2.1.2(47) using the parameters ‘ --format BAMPE --gsize hs --keep-dup 1 --bdg --SPMR --qvalue 0.05’. Candidate regions were further filtered by fold enrichment score. Differential binding analysis was run on peaks which were found in at least 2 samples using DiffBind v2.10.0 on R v4.3.1 (http://bioconductor.org/packages/release/bioc/vignettes/DiffBind/inst/doc/DiffBind.pdf(48) and R Core Team 2023). The input was the binding region coordinates and associated q-values from MACS2, and mapped reads for each sample. Differential binding analysis was performed based on the comparison of KS1 versus control samples with Benjamini-Hochberg adjusted p-value threshold of 0.05. Bedtools(49) intersect tools was used to intersect enhancer regions from EpiMap repository(50) based on iPSC DF19.11, BSS01370 Neural Progenitor or BSS01366 Neural cells with H3K4me1 regions from ChIP-Seq analysis with minimum intersect ratio of 0.5 in relation to enhancer region. Deeptools v3.4.3(51) was used to compare the distribution and intensity of H3K4me1 peaks across samples in different genomic regions. BamCompare was used to subtract ChIP input reads from each sample, whilst being simultaneously normalised to bins per million (BPM) with bin size of 20. ComputeMatrix created a matrix of reads for the normalised reads of each sample relative to the sites of interest used for visualisation, which was then passed to plotHeatmap for visualisation of read distribution within regions of interest. HOMER software was used for annotation of peaks(52). Analysis of ChIP-seq data between different cell time points was performed using ChIPpeakAnno v3.36.0 package within R v4.3.1.

For motif analysis, unique peak summits from samples within differential H3K4me1 regions were extracted using Bedtools(49) intersect. Flanking regions of ±250bp were added to the peak summits (Bedtools(49) slop) and fasta sequences within those regions were extracted using Bedtools(49) getfasta. MEME-TomTom within MEME-ChIP was used to search JASPAR CORE(53) vertebrates, UniProbe Mouse(54) and Jolma 2013 Human and Mouse databases(55) for enriched motifs within differential H3K4me1 regions.

### Statistical analysis

Two-tailed T-test was performed to analyse the differences in mean expression values of KMT2D and intensity of H3K4me1/2/3. Hypergeometric test was performed to assess over-representation of DEGs per chromosome and per chromosomal bins. Two-tailed Fisher’s test was performed to analyse the association of H3K4me1 loss within the enhancer regions. Spearman correlation test was performed to compare correlation of Log2 Fold Changes of DEGs from Dataset 1 Neural Progenitors and Log 2 Fold Changes calculated from Dataset 3 Neural Progenitors.

## ACKNOWLEGMENTS

We thank KS1 individuals and their families for providing the samples. We acknowledge the support of Great Ormond Street Hospital Charity (V4621), Newlife Charity (Grant reference 16-17/10), Manchester University Hospitals NHS Foundation Trust Kabuki Research Fund no. 629396, the NIHR Manchester Biomedical Research Centre (NIHR203308), and the MRC Epigenomics of Rare Diseases Node (MR/Y008170/1). We also thank Julieta O’Flaherty and Steven Woods for the control iPSCs (JF191b(36), SW171a(37)) and the Genomic Technologies Core Facility at the University of Manchester for RNA and ChIP library preparation and sequencing. This study makes use of control and KS1 iPSCs and RNAseq data generated by the HipSci consortium (www.hipsci.org), funded by The Wellcome Trust and the MRC.

## REFERENCES

1. Froimchuk E, Jang Y, Ge K. Histone H3 lysine 4 methyltransferase KMT2D. Gene. 2017;627:337–42.

2. Shilatifard A. The COMPASS family of histone H3K4 methylases: mechanisms of regulation in development and disease pathogenesis. Annu Rev Biochem. 2012;81:65–95.

3. Lee JE, Wang C, Xu S, Cho YW, Wang L, Feng X, et al. H3K4 mono- and di-methyltransferase MLL4 is required for enhancer activation during cell differentiation. Elife. 2013;2:e01503.

4. Wang C, Lee JE, Lai B, Macfarlan TS, Xu S, Zhuang L, et al. Enhancer priming by H3K4 methyltransferase MLL4 controls cell fate transition. Proc Natl Acad Sci U S A. 2016;113(42):11871–6.

5. Cao K, Collings CK, Morgan MA, Marshall SA, Rendleman EJ, Ozark PA, et al. An Mll4/COMPASS-Lsd1 epigenetic axis governs enhancer function and pluripotency transition in embryonic stem cells. Sci Adv. 2018;4(1):eaap8747.

6. Van Laarhoven PM, Neitzel LR, Quintana AM, Geiger EA, Zackai EH, Clouthier DE, et al. Kabuki syndrome genes KMT2D and KDM6A: functional analyses demonstrate critical roles in craniofacial, heart and brain development. Hum Mol Genet. 2015;24(15):4443–53.

7. Bjornsson HT, Benjamin JS, Zhang L, Weissman J, Gerber EE, Chen YC, et al. Histone deacetylase inhibition rescues structural and functional brain deficits in a mouse model of Kabuki syndrome. Sci Transl Med. 2014;6(256):256ra135.

8. Ng SB, Bigham AW, Buckingham KJ, Hannibal MC, McMillin MJ, Gildersleeve HI, et al. Exome sequencing identifies MLL2 mutations as a cause of Kabuki syndrome. Nat Genet. 2010;42(9):790–3.

9. Banka S, Veeramachaneni R, Reardon W, Howard E, Bunstone S, Ragge N, et al. How genetically heterogeneous is Kabuki syndrome?: MLL2 testing in 116 patients, review and analyses of mutation and phenotypic spectrum. Eur J Hum Genet. 2012;20(4):381–8.

10. Banka S, Lederer D, Benoit V, Jenkins E, Howard E, Bunstone S, et al. Novel KDM6A (UTX) mutations and a clinical and molecular review of the X-linked Kabuki syndrome (KS2). Clin Genet. 2015;87(3):252–8.

11. Miyake N, Koshimizu E, Okamoto N, Mizuno S, Ogata T, Nagai T, et al. MLL2 and KDM6A mutations in patients with Kabuki syndrome. Am J Med Genet A. 2013;161A(9):2234–43.

12. Adam MP, Banka S, Bjornsson HT, Bodamer O, Chudley AE, Harris J, et al. Kabuki syndrome: international consensus diagnostic criteria. J Med Genet. 2019;56(2):89–95.

13. Fasciani A, D’Annunzio S, Poli V, Fagnocchi L, Beyes S, Michelatti D, et al. MLL4-associated condensates counterbalance Polycomb-mediated nuclear mechanical stress in Kabuki syndrome. Nat Genet. 2020;52(12):1397–411.

14. Jung YL, Hung C, Choi J, Lee EA, Bodamer O. Characterizing the molecular impact of KMT2D variants on the epigenetic and transcriptional landscapes in Kabuki syndrome. Hum Mol Genet. 2023;32(13):2251–61.

15. Carosso GA, Boukas L, Augustin JJ, Nguyen HN, Winer BL, Cannon GH, et al. Precocious neuronal differentiation and disrupted oxygen responses in Kabuki syndrome. JCI Insight. 2019;4(20).

16. Jefri M, Zhang X, Stumpf PS, Zhang L, Peng H, Hettige N, et al. Kabuki syndrome stem cell models reveal locus specificity of histone methyltransferase 2D (KMT2D/MLL4). Hum Mol Genet. 2022;31(21):3715–28.

17. Faundes V, Malone G, Newman WG, Banka S. A comparative analysis of KMT2D missense variants in Kabuki syndrome, cancers and the general population. J Hum Genet. 2019;64(2):161–70.

18. Faundes V, Goh S, Akilapa R, Bezuidenhout H, Bjornsson HT, Bradley L, et al. Clinical delineation, sex differences, and genotype-phenotype correlation in pathogenic KDM6A variants causing X-linked Kabuki syndrome type 2. Genet Med. 2021;23(7):1202–10.

19. Shi Y, Kirwan P, Livesey FJ. Directed differentiation of human pluripotent stem cells to cerebral cortex neurons and neural networks. Nat Protoc. 2012;7(10):1836–46.

20. El-Brolosy MA, Kontarakis Z, Rossi A, Kuenne C, Gunther S, Fukuda N, et al. Genetic compensation triggered by mutant mRNA degradation. Nature. 2019;568(7751):193-7.

21. Cocciadiferro D, Augello B, De Nittis P, Zhang J, Mandriani B, Malerba N, et al. Dissecting KMT2D missense mutations in Kabuki syndrome patients. Hum Mol Genet. 2018;27(21):3651–68.

22. McConnell BB, Yang VW. Mammalian Kruppel-like factors in health and diseases. Physiol Rev. 2010;90(4):1337–81.

23. Cassandri M, Smirnov A, Novelli F, Pitolli C, Agostini M, Malewicz M, et al. Zinc-finger proteins in health and disease. Cell Death Discov. 2017;3:17071.

24. Guo JK, Menke AL, Gubler MC, Clarke AR, Harrison D, Hammes A, et al. WT1 is a key regulator of podocyte function: reduced expression levels cause crescentic glomerulonephritis and mesangial sclerosis. Hum Mol Genet. 2002;11(6):651–9.

25. Giguere V. Transcriptional control of energy homeostasis by the estrogen-related receptors. Endocr Rev. 2008;29(6):677–96.

26. Mousavi Khorshidi MS, Seeleuthner Y, Chavoshzadeh Z, Behfar M, Hamidieh AA, Alimadadi H, et al. Clinical, Immunological, and Genetic Findings in Iranian Patients with MHC-II Deficiency: Confirmation of c.162delG RFXANK Founder Mutation in the Iranian Population. J Clin Immunol. 2023;43(8):1941–52.

27. Kuehn HS, Boast B, Rosenzweig SD. Inborn errors of human IKAROS: LOF and GOF variants associated with primary immunodeficiency. Clin Exp Immunol. 2023;212(2):129–36.

28. Larsen KB, Lutterodt MC, Mollgard K, Moller M. Expression of the homeobox genes OTX2 and OTX1 in the early developing human brain. J Histochem Cytochem. 2010;58(7):669–78.

29. Kashyap V, Laursen KB, Brenet F, Viale AJ, Scandura JM, Gudas LJ. RARgamma is essential for retinoic acid induced chromatin remodeling and transcriptional activation in embryonic stem cells. J Cell Sci. 2013;126(Pt 4):999–1008.

30. Fontanet PA, Rios AS, Alsina FC, Paratcha G, Ledda F. Pea3 Transcription Factors, Etv4 and Etv5, Are Required for Proper Hippocampal Dendrite Development and Plasticity. Cereb Cortex. 2018;28(1):236–49.

31. Kim M, Jang YJ, Lee M, Guo Q, Son AJ, Kakkad NA, et al. The transcriptional regulatory network modulating human trophoblast stem cells to extravillous trophoblast differentiation. Nat Commun. 2024;15(1):1285.

32. Choi HJ, Geng Y, Cho H, Li S, Giri PK, Felio K, et al. Differential requirements for the Ets transcription factor Elf-1 in the development of NKT cells and NK cells. Blood. 2011;117(6):1880–7.

33. Kakun RR, Melamed Z, Perets R. PAX8 in the Junction between Development and Tumorigenesis. Int J Mol Sci. 2022;23(13).

34. Thompson JA, Ziman M. Pax genes during neural development and their potential role in neuroregeneration. Prog Neurobiol. 2011;95(3):334–51.

35. Loh CH, Veenstra GJC. The Role of Polycomb Proteins in Cell Lineage Commitment and Embryonic Development. Epigenomes. 2022;6(3).

36. Mancini FE, Humphreys PEA, Woods S, Bates N, Cuvertino S, O’Flaherty J, et al. Effect of a retinoic acid analogue on BMP-driven pluripotent stem cell chondrogenesis. 2023.

37. Wood KA, Rowlands CF, Thomas HB, Woods S, O’Flaherty J, Douzgou S, et al. Modelling the developmental spliceosomal craniofacial disorder Burn-McKeown syndrome using induced pluripotent stem cells. PLoS One. 2020;15(7):e0233582.

38. Bolger AM, Lohse M, Usadel B. Trimmomatic: a flexible trimmer for Illumina sequence data. Bioinformatics. 2014;30(15):2114–20.

39. Dobin A, Davis CA, Schlesinger F, Drenkow J, Zaleski C, Jha S, et al. STAR: ultrafast universal RNA-seq aligner. Bioinformatics. 2013;29(1):15–21.

40. Danecek P, Bonfield JK, Liddle J, Marshall J, Ohan V, Pollard MO, et al. Twelve years of SAMtools and BCFtools. Gigascience. 2021;10(2).

41. Liao Y, Smyth GK, Shi W. featureCounts: an efficient general purpose program for assigning sequence reads to genomic features. Bioinformatics. 2014;30(7):923–30.

42. Kuleshov MV, Jones MR, Rouillard AD, Fernandez NF, Duan Q, Wang Z, et al. Enrichr: a comprehensive gene set enrichment analysis web server 2016 update. Nucleic Acids Res. 2016;44(W1):W90–7.

43. Chen EY, Tan CM, Kou Y, Duan Q, Wang Z, Meirelles GV, et al. Enrichr: interactive and collaborative HTML5 gene list enrichment analysis tool. BMC Bioinformatics. 2013;14:128.

44. Briggs P, Hunter AL, Yang SH, Sharrocks AD, Iqbal M. PEGS: An efficient tool for gene set enrichment within defined sets of genomic intervals. F1000Res. 2021;10:570.

45. Wingett SW, Andrews S. FastQ Screen: A tool for multi-genome mapping and quality control. F1000Res. 2018;7:1338.

46. Langmead B, Salzberg SL. Fast gapped-read alignment with Bowtie 2. Nat Methods. 2012;9(4):357–9.

47. Zhang Y, Liu T, Meyer CA, Eeckhoute J, Johnson DS, Bernstein BE, et al. Model-based analysis of ChIP-Seq (MACS). Genome Biol. 2008;9(9):R137.

48. Ross-Innes CS, Stark R, Teschendorff AE, Holmes KA, Ali HR, Dunning MJ, et al. Differential oestrogen receptor binding is associated with clinical outcome in breast cancer. Nature. 2012;481(7381):389-93.

49. Quinlan AR, Hall IM. BEDTools: a flexible suite of utilities for comparing genomic features. Bioinformatics. 2010;26(6):841–2.

50. Boix CA, James BT, Park YP, Meuleman W, Kellis M. Regulatory genomic circuitry of human disease loci by integrative epigenomics. Nature. 2021;590(7845):300-7.

51. Ramirez F, Ryan DP, Gruning B, Bhardwaj V, Kilpert F, Richter AS, et al. deepTools2: a next generation web server for deep-sequencing data analysis. Nucleic Acids Res. 2016;44(W1):W160–5.

52. Heinz S, Benner C, Spann N, Bertolino E, Lin YC, Laslo P, et al. Simple combinations of lineage-determining transcription factors prime cis-regulatory elements required for macrophage and B cell identities. Mol Cell. 2010;38(4):576–89.

53. Castro-Mondragon JA, Riudavets-Puig R, Rauluseviciute I, Lemma RB, Turchi L, Blanc-Mathieu R, et al. JASPAR 2022: the 9th release of the open-access database of transcription factor binding profiles. Nucleic Acids Res. 2022;50(D1):D165–D73.

54. Hume MA, Barrera LA, Gisselbrecht SS, Bulyk ML. UniPROBE, update 2015: new tools and content for the online database of protein-binding microarray data on protein-DNA interactions. Nucleic Acids Res. 2015;43(Database issue):D117–22.

55. Jolma A, Yan J, Whitington T, Toivonen J, Nitta KR, Rastas P, et al. DNA-binding specificities of human transcription factors. Cell. 2013;152(1-2):327–39.

