## Supplemental Table 1 and Supplemental Figures S1-S11 for "Epigenome and transcriptome changes in *KMT2D*-related Kabuki syndrome Type 1 iPSCs, neuronal progenitors and cortical neurons"

| | Controls | | | Kabuki syndrome 1 | | | | | | | |
|---|---|---|---|---|---|---|---|---|---|---|---|
| | **# Samples/Sample name** | **Sex** | **Tissue** | **Sample name** | **Mutation** | **Mutation type** | **# Samples** | **Sex** | **Tissue** | **RNAseq** | **ChIPseq** |
| **Dataset 1 (DS1)** | 3 | M | DSFC | iPSC 3.1/3.2/3.3 | c.16019G>A p.(Arg5340Gln) | missense | 3 | F | DSFC | x | x |
| | | 42F | DSFC | iPSC 1 | c.5527dupA | frameshift | 1 | F | DSFC | x | |
| **Dataset 2 (DS2)** | 50 | | | iPSC 2.1/2.2 | c.16180G>T p.(Glu5394Ter) | nonsense | 2 | M | DSFC | x | |
| | | 8M | DSFC | iPSC 3 | c.16019G>A p.(Arg5340Gln) | missense | 1 | F | DSFC | x | |
| | Ctrl 1 | M | DSFC | iPSC 1 | c.5527dupA | frameshift | 1 | F | DSFC | x | |
| **Dataset 3 (DS3)** | Ctrl 2 | F | PBMC | iPSC 2 | c.16180G>T p.(Glu5394Ter) | nonsense | 1 | M | DSFC | x | |
| | Ctrl 3 | M | DSFC | iPSC 3 | c.16019G>A p.(Arg5340Gln) | missense | 1 | F | DSFC | x | |

**Supplementary Table 1. List of samples for the integrated RNAseq datasets.** Samples number, name, mutation and sex are listed. DSFC = dermal skin fibroblast cells; PBMC = peripheral blood mononuclear cells.

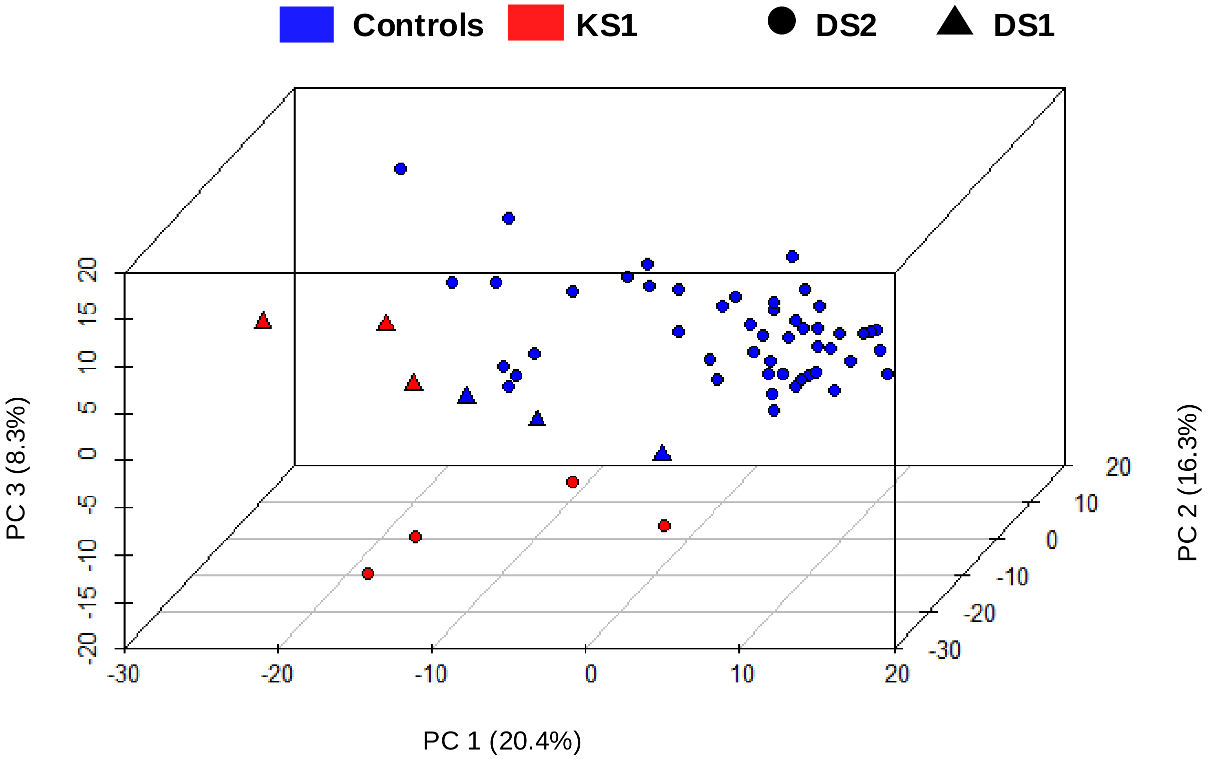


**Supplementary Figure 1. Integration of RNAseq datasets.** PCA plot showing distinct clustering of the two groups (control: blue; KS1: red).

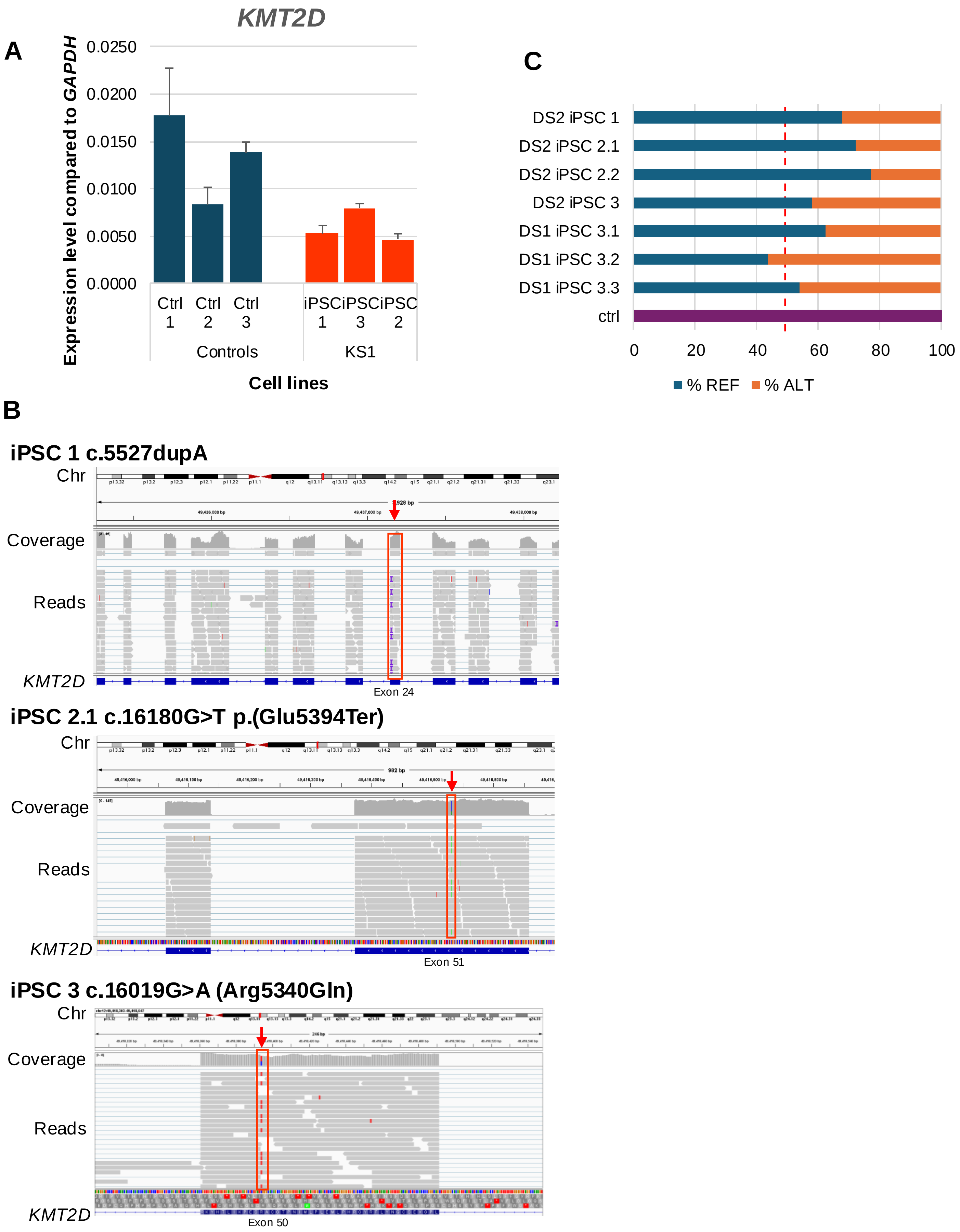


**Supplementary Figure 2. Analysis of *KMT2D* expression A)** qRT-PCR analysis of *KMT2D* transcript level relative to *GAPDH* in control and KS1 iPSCs (frameshift c.5527dupA, missense (c.16019G>A, p.(Arg5340Gln)), and nonsense (c.16180G>T, p.(Glu5394Ter)) variants (n=3)). **B)** Mutant read visualization using IGV. **C)** Bar graph showing percentage of REF (reads with reference – no mutation; blue for control and light blue for KS1) and ALT (reads with mutation, orange) (DS2 iPSC1 c.5527dupA, DS2 iPSC 2.1 c.16180G>T, DS2 iPSC 2.2 c.16180G>T, DS2 iPSC 3 c.16019G>A, DS1 iPSC 3.1 c.16019G>A, DS1 iPSC 3.2 c.16019G>A, DS1 iPSC 3.3 c.16019G>A).

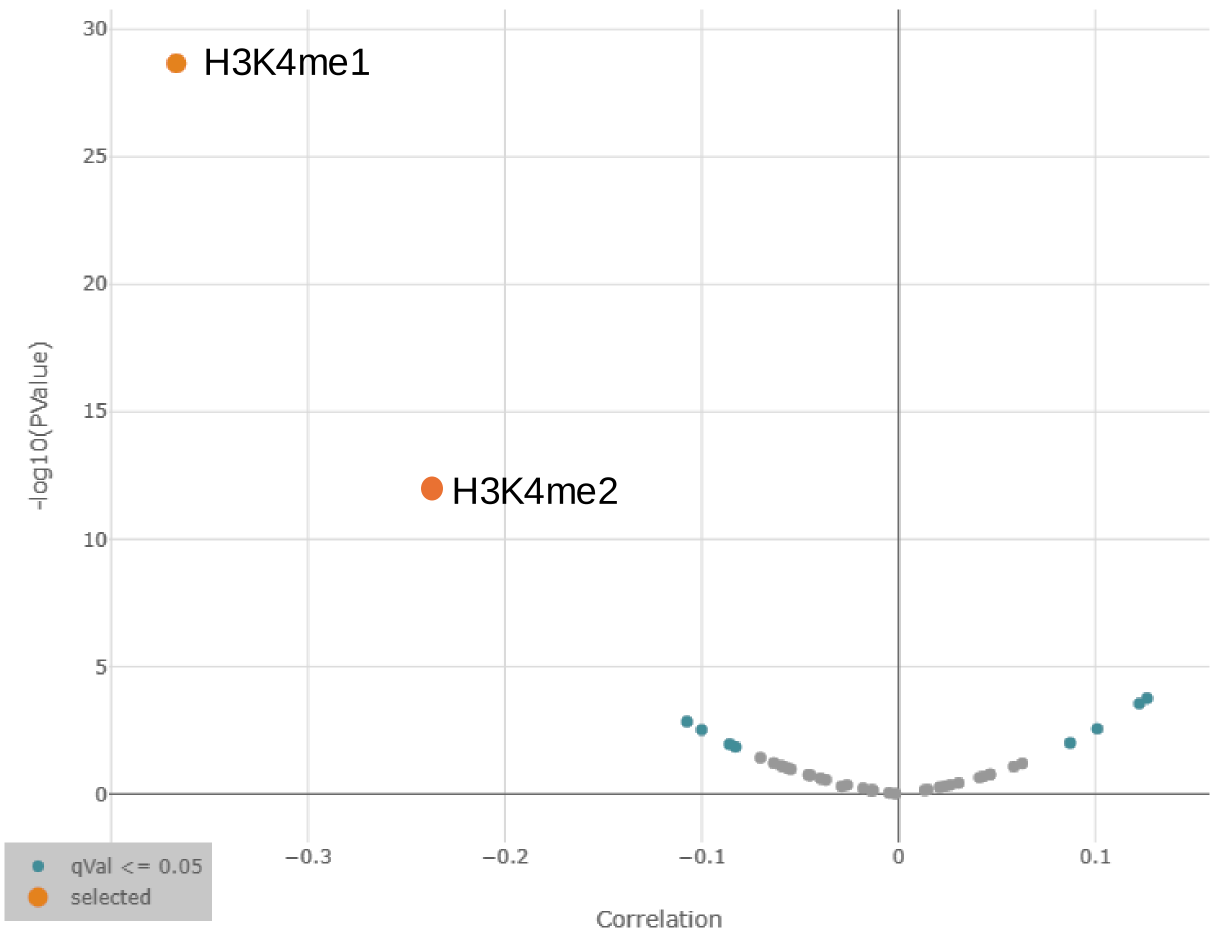


**Supplementary Figure 3. Analysis of H3K4 methylation using DepMap.** Volcano plot showing the level of histone marks in cell lines carrying *KMT2D* mutations using DepMap portal (https://depmap.org/portal). H3K4me1 and H3K4me2 levels are significantly reduced.

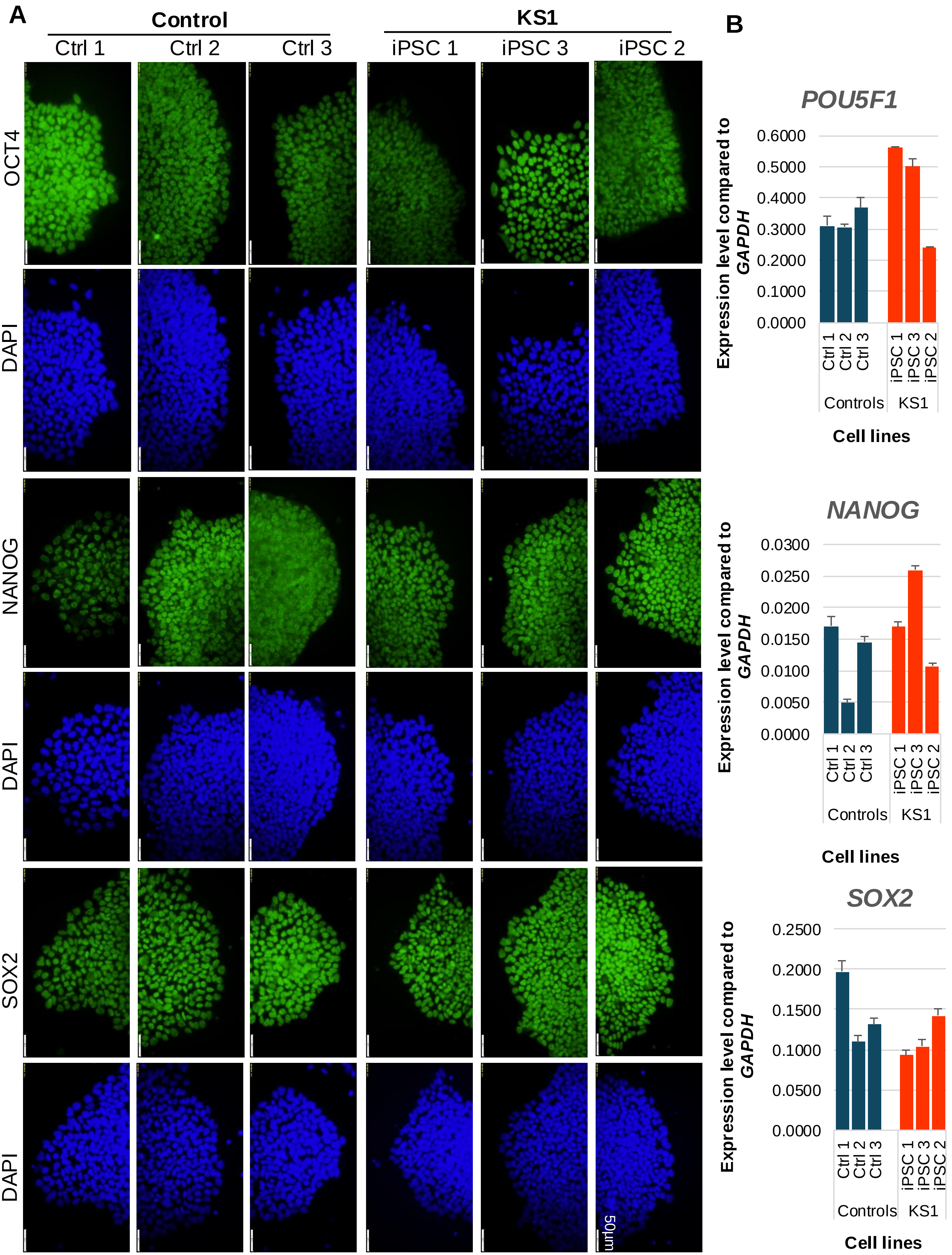


**Supplementary Figure 4. Validation analysis for pluripotency in KS1 and control iPSC lines. A)** Immunofluorescence staining for OCT4, NANOG and SOX2 in iPSCs. **B)** qRT-PCR analysis of *POU5F1*, *NANOG* and *SOX2* transcript level relative to *GAPDH* in control (Ctrl 1, Ctrl 2, Ctrl 3) and KS1 iPSCs (iPSC 1, iPSC 3, iPSC 2) (n=3).

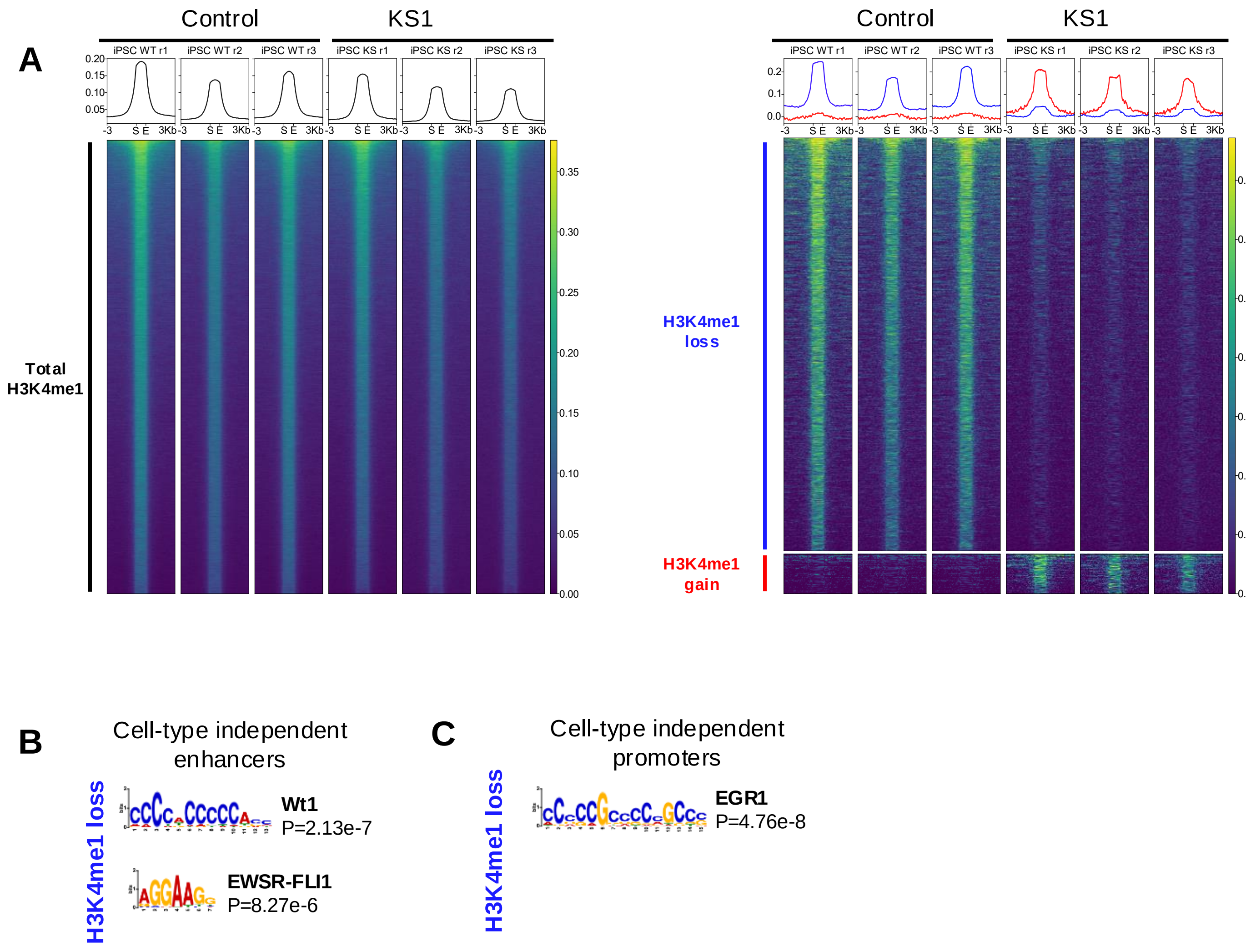


**Supplementary Figure 5. Analysis of H3K4me1 level in different genomic regions in iPSCs. A)** Heatmaps of H3K4me1 regions mapped to total (n=81,930 in black; left), H3K4me1 loss (n=3647 in blue; right) and gain (n=349 in red; right) EpiMap enhancer regions based on all cell-types. Top TF motifs predicted in differential H3K4me1 peaks (loss and gain of H3K4me1 in binding regions) between KS1 and control in **B)** cell–type independent enhancers and **C)** cell-type independent promoters.

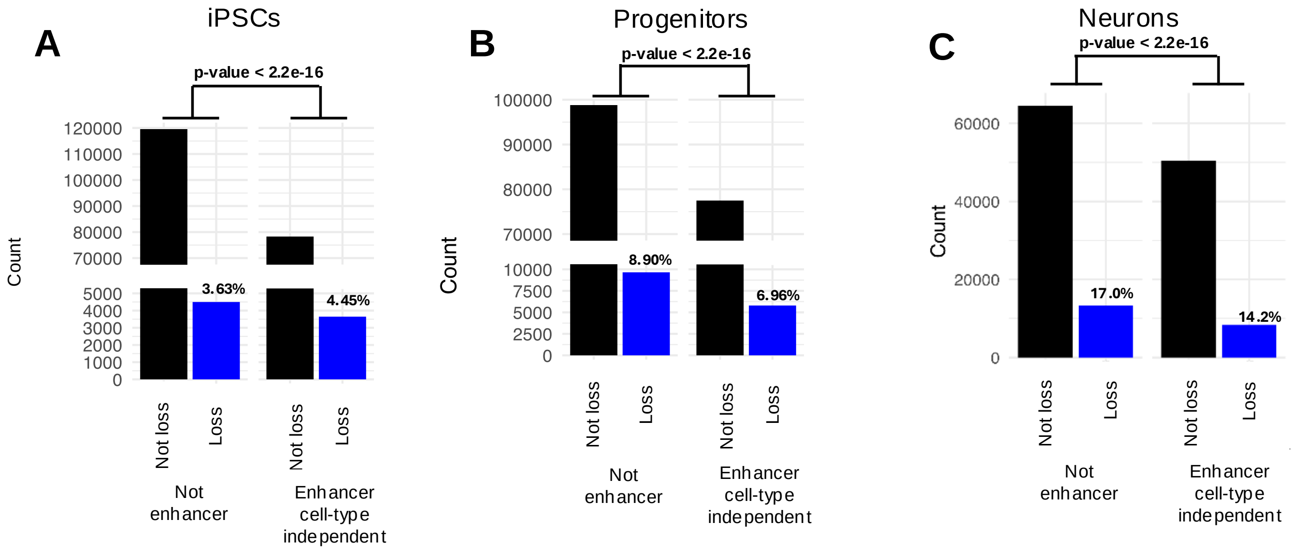


**Supplementary Figure 6. Association analysis of loss of H3K4me1 in KS1 iPSC, progenitors and neurons.** Analysis of association of H3K4me1 loss with enhancer regions at day 0 iPSC stage **(A)**, at day 18 neuronal progenitor stage **(B)** and at day 30 neuronal stage **(C)** (two-tailed fisher's exact test) with percentage of loss per region.

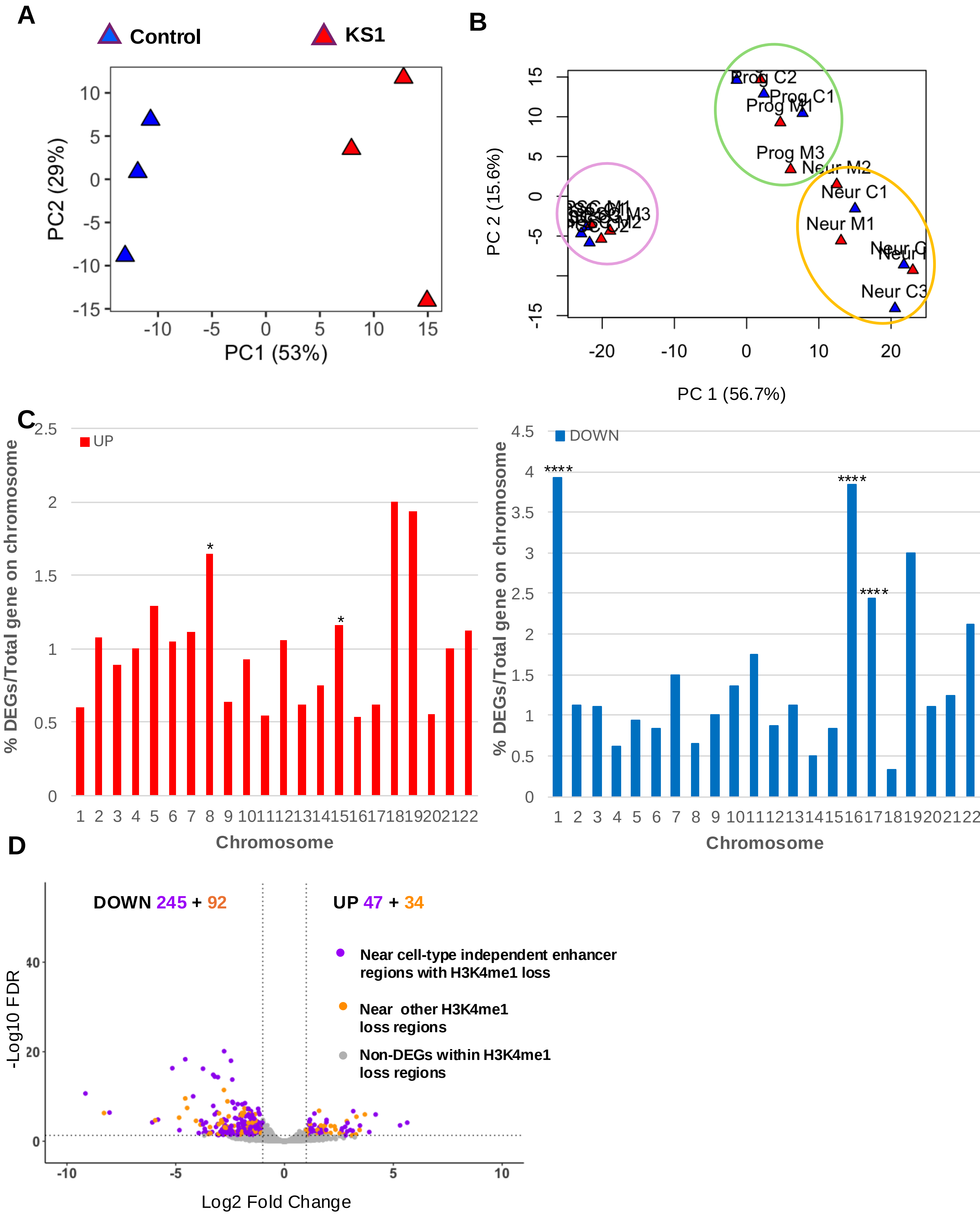


**Supplementary Figure 7. Analysis of the transcriptome in KS1 iPSCs. A)** PCA plot showing distinct clustering of the two groups (control: blue; KS1 (c.16019G>A): red) at the iPSC stage in DS1. **B)** PCA plot showing all the three cell stage Blue circle show iPSC, green progenitors and orange neuronal cells (control: blue; KS1: red). **C)** Bar graph showing percentage of DEGs per total gene on chromosome in KS1 compared to controls. Upregulated genes are in red and downregulated genes are in blue. Hypergeometric test was performed to assess the enrichment of DEGs in each chromosome (*FDR < 0.05; ****FDR < 0.0001; hypergeometric test). **D)** Volcano plot showing DEGs within ±100kb of H3K4me1 loss regions.

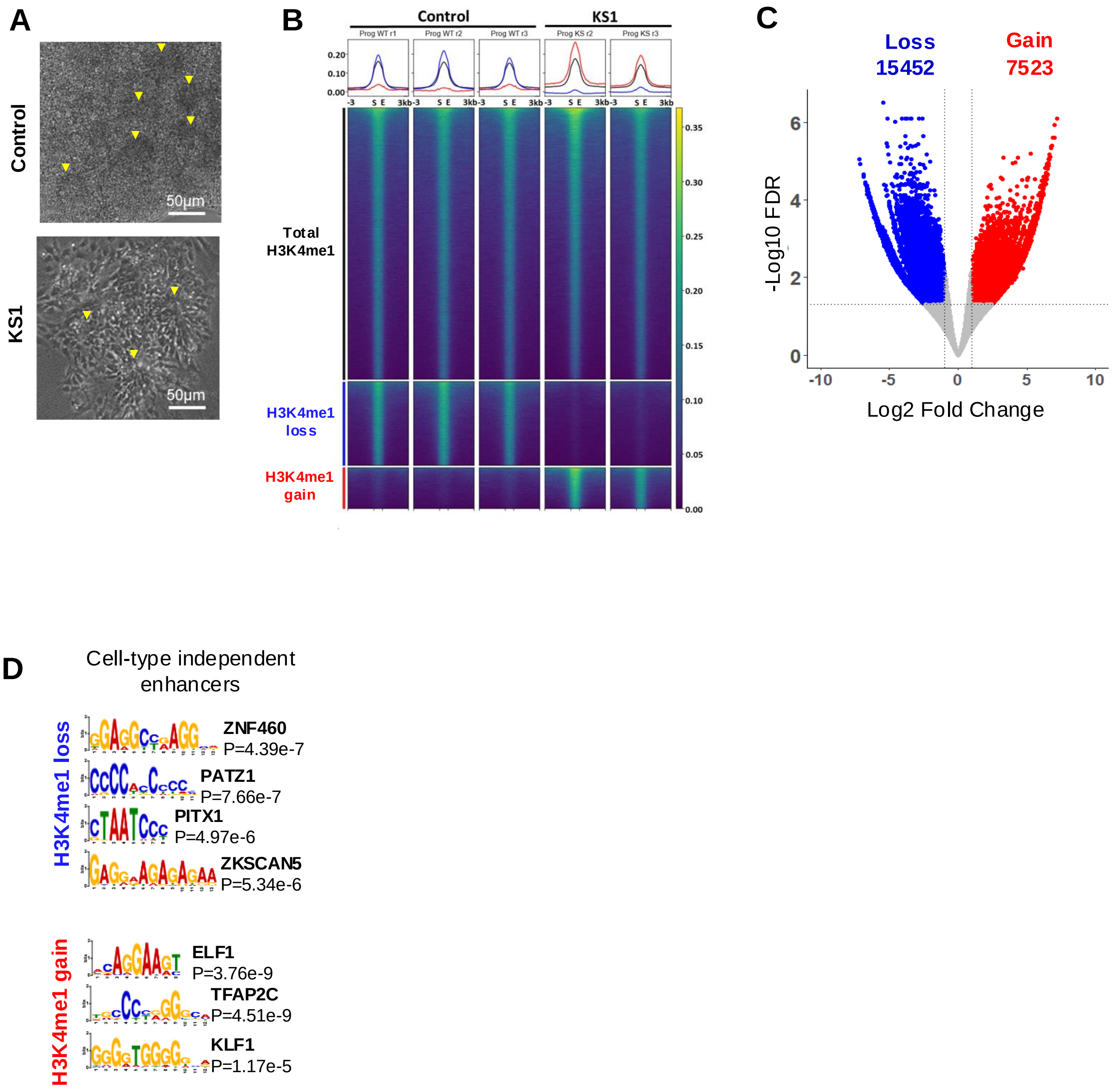


**Supplementary Figure 8. Analysis of H3K4me1 level in KS1 neuronal progenitor cells. A)** Representative phase contrast images of control and KS1 neural progenitor (Day 18) morphology. Rosettes are highlighted by yellow arrowheads. **B)** Heatmap of H3K4me1 peaks (black), H3K4me1 loss regions (blue), and H3K4me1 gain regions (red) for neuronal progenitors. **C)** Volcano plot showing loss (blue) and gain (red) in H3K4me1. **D)** Top TF motifs predicted in differential H3K4me1 peaks (loss and gain of H3K4me1 in binding regions) between KS1 and control in cell–type independent enhancers.

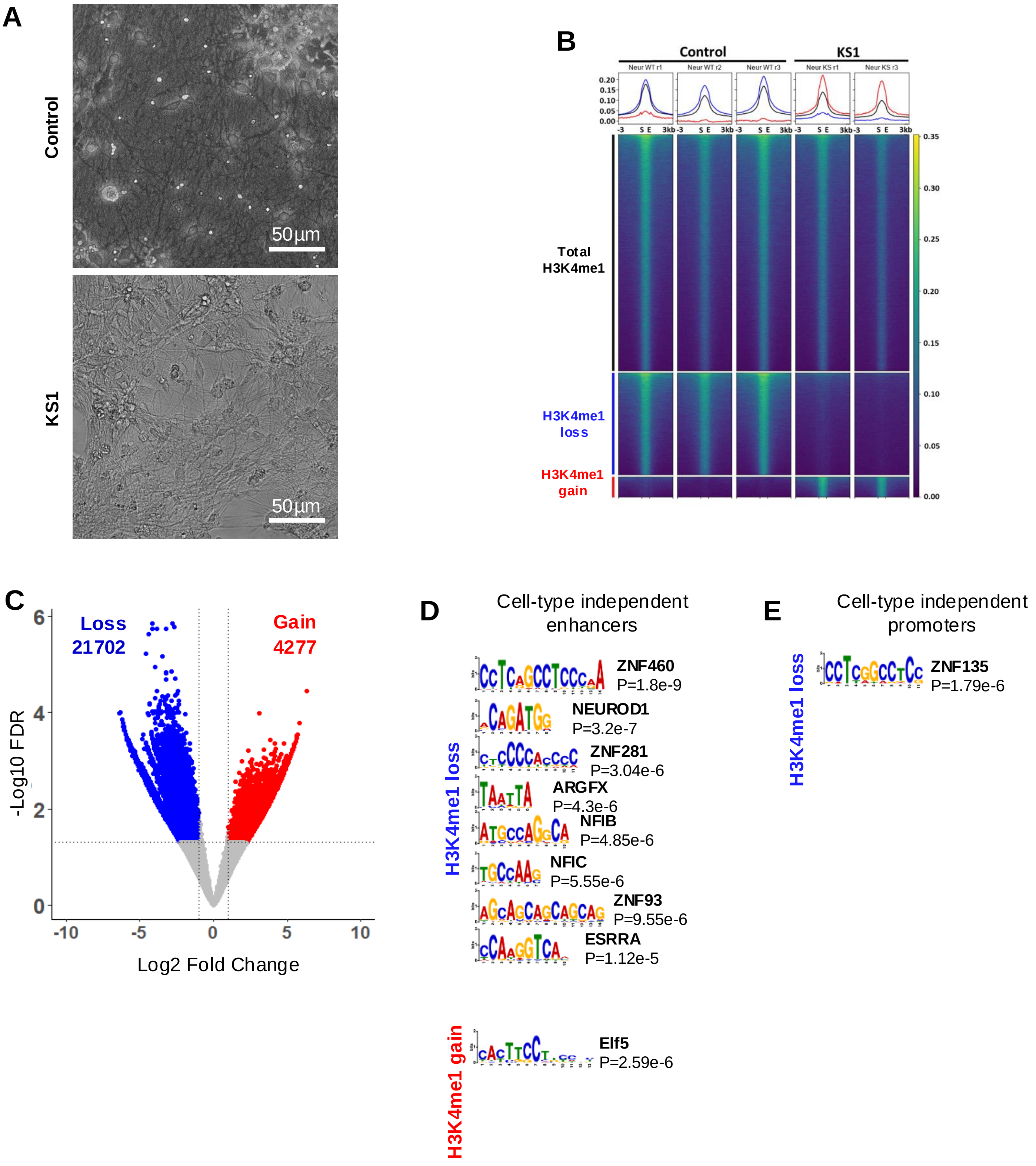


**Supplementary Figure 9. Analysis of H3K4me1 level in different genomic regions in neuronal cells. A)** Representative phase contrast images of control and KS1 neuronal morphology **B)** Heatmap of H3K4me1 peaks (black), H3K4me1 loss regions (blue), and H3K4me1 gain regions (red) for neurons. **C)** Volcano plot showing loss (blue) and gain (red) in H3K4me1. Top TF motifs predicted in differential H3K4me1 peaks (loss and gain of H3K4me1 in binding regions) between KS1 and control in **D)** cell–type independent enhancers and **E)** cell-type independent promoters.

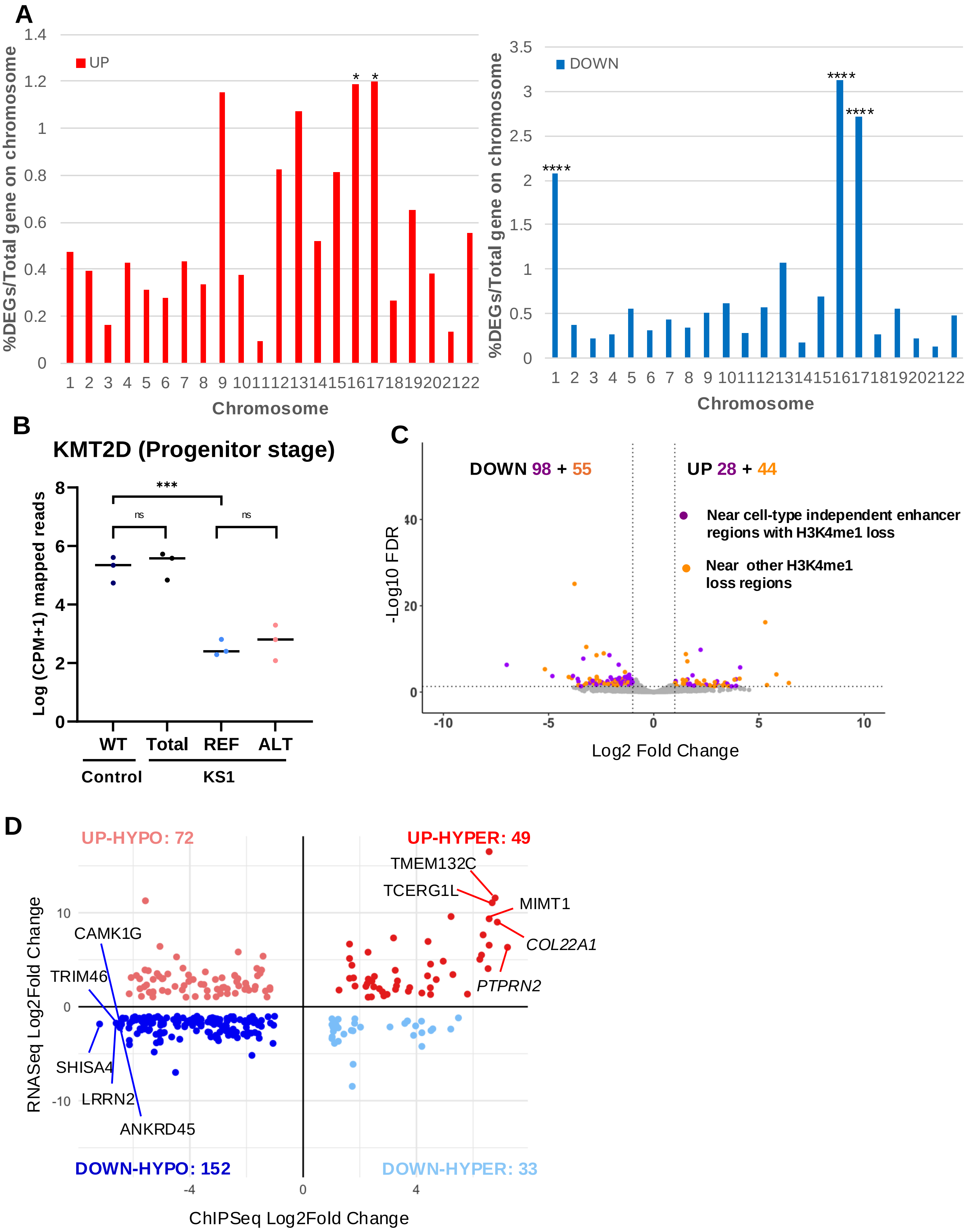


**Supplementary Figure 10. Analysis of the transcriptome in KS1 neuronal progenitor cells. A)** Bar graph showing percentage of DEGs per total gene on chromosome in KS1 compared to controls. Upregulated genes are in red and downregulated genes are in blue. Hypergeometric test was performed to assess the enrichment of DEGs in each chromosome (*FDR<0.05; ****FDR < 0.0001; hypergeometric test). **B)** Volcano plot showing down and up DEGs within ±100kb of differentially lost H3K4me1 regions. **C)** Scatter plot showing change in expression and change in H3K4me1 within ±100kb of DEGs.

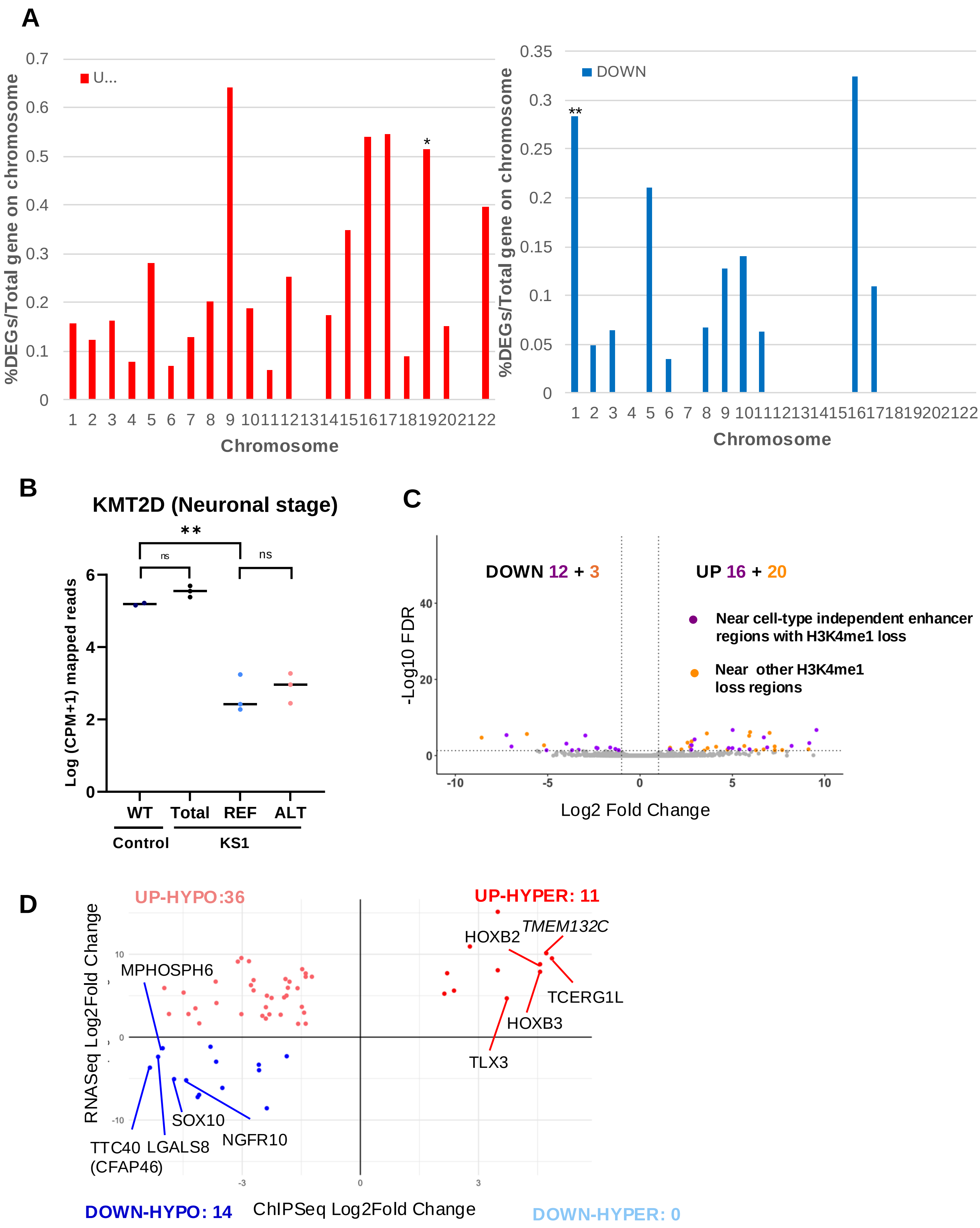


**Supplementary Figure 11. Analysis of the transcriptome in KS1 neuronal cells. A)** Bar graph showing percentage of DEGs per total gene on chromosome in KS1 compared to controls. Upregulated genes are in red and downregulated genes are in blue. Hypergeometric test was performed to assess the enrichment of DEGs in each chromosome (*FDR<0.05; **FDR < 0.01; hypergeometric test). **B)** Volcano plot showing down and up DEGs within ±100kb of differentially lost H3K4me1 regions. **C)** Scatter plot showing change in expression and change in H3K4me1 within ±100kb of DEGs.
